# Characterizing spatial functional microniches with SpaceTravLR

**DOI:** 10.1101/2025.11.13.688264

**Authors:** Koushul Ramjattun, Alyson Wang, Hannah Lee, Shilpi Giri, Yijia Chen, William A. MacDonald, Nathan D. Lord, Amanda C. Poholek, Youjin Lee, Jishnu Das

**Author notes:** co-first author/equal contribution (Koushul Ramjattun, Alyson Wang). Corresponding authors – Jishnu Das, Youjin Lee, Amanda Poholek.

## Abstract

The advent of spatial omics has revolutionized our understanding of tissue biology; however, these technologies remain largely descriptive and do not capture how changes in gene regulation propagate across spatial neighborhoods. While *in-silico* perturbation methods and foundation models aim to model the impact of genetic perturbations, these methods are limited to single-cell approaches that lack spatial resolution. Other studies can delineate morphological domains based on transcriptional similarity, but not spatial functional microniches. We address this major unmet need by developing SpaceTravLR (Spatially perturbing Transcription factors, Ligands and Receptors), a novel interpretable machine learning approach that generalizes across tissues and species, uncovering spatial features linked to functional outcomes, thereby capturing functional microniches with spatial resolution. SpaceTravLR infers how single or combinatorial genetic perturbations rewire signals across the tissue neighborhood, by propagating effects through underlying spatially resolved molecular networks, thereby modeling how perturbations can reshape both the targeted cell and its surrounding neighborhood. SpaceTravLR defines novel spatial microniches across a range of tissues at different scales of organization (niches, neighborhoods and tissues), disease and developmental contexts. SpaceTravLR’s perturbation predictions are made solely from spatial omics data and closely align with experimental validation or known outcomes based on mechanistic studies. Critically, our approach enables the generation of mechanistic hypotheses underlying identified niches. We show SpaceTravLR discovered a novel mechanism for *Ccr4* that drives the spatial location of a pathogenic population of allergen-specific T helper 2 (Th2) cells as they develop in the lymph node, which was experimentally validated in a murine model. Overall, SpaceTravLR provides a novel interpretable and experimentally validated framework for uncovering how genes act individually and combinatorially through cell-intrinsic and cell-extrinsic circuits to shape spatial tissue organization and function.

## Main

Cells can influence one another through a network of secreted proteins and cell surface molecules to extend their transcriptional programs beyond their own cytoplasm and thus functionally impact the states and fates of neighboring cells. A holistic understanding of tissue biology and organization requires connecting each cell’s molecular activity to its spatial context within the tissue. Combined with genome perturbation tools such as CRISPR^1^, these advances hold enormous promise towards identifying causal modulators driving functional phenotypes, particularly when paired with high throughput single-cell RNA sequencing (scRNA-seq). Emerging technologies such as Perturb-seq^2^ and CROP-seq^3^ have already shown great potential for causal gene modulatory network reconstruction. However, these methods fail to recover the spatial context within which a perturbation occurs. To gain a holistic understanding of tissue biology and organization, we must link the spatial context of a cell to its biology using tools that capture interacting transcriptomes and perturb genes to identify causal drivers of function.

Here, we introduce SpaceTravLR, a first-in-class framework to advance spatial biology at single cell resolution from descriptive mapping to causal modeling by uncovering functional microniches directly from spatial transcriptomics (ST) data. SpaceTravLR connects the downstream effect of gene perturbation to the spatial neighborhoods in which those effects occur, thereby linking molecular function to local tissue organization. The integration of *in-silico* perturbation screening with spatial context allows us to infer how the combination of gene-gene interactions rewire signaling to define distinct cellular environments.

We demonstrate the capabilities of SpaceTravLR by applying it to well-studied systems and accurately recapitulating perturbation outcomes across T and B cell populations in the germinal center, that are supported by mechanistic experiments. We further highlight SpaceTravLR’s unique ability to infer functional cell-cell communication using a new true single cell spatial murine kidney dataset, where we identify microniches associated with distinct macrophage transition outcomes based on expression of the ligand *Mif* in surrounding epithelial cells. We validate our model in the context of embryo development using a spatially profiled knockout dataset, showing accurate predictions for unseen cell types and neural structures that arise in Tbx6 mutants. To screen for novel gene candidates that would manifest desired cellular fates, we systematically perturbed all transcription factors (TF), ligands and receptors across all cell types in human melanoma and a murine model of allergic asthma. In melanoma, we predict enhanced CD8 T cell anti-tumor activity from specific TF knockouts. Finally, in the murine asthma model, we predict and experimentally validate an unconventional migration of developing allergen-specific pathogenic T helper 2 (Th2) cells in the lung draining lymph node from the T cell zone to the B cell zone following disrupted *Ccr4* signaling.

### An interpretable machine learning model for discovering functional microniches from spatial transcriptomics data

SpaceTravLR aims to push spatial analyses at single cell resolution from descriptive correlations toward functional mechanistic insights. To this end, we leverage ST data to uncover tissue regions where local differences in the cellular microenvironment drive divergent cell fates in neighboring cells within tissues (**Fig. 1A**). SpaceTravLR simulates how gene perturbations affect both the perturbed cell and its spatial neighbors. The model learns the magnitude and direction of impact that perturbing one gene has on downstream target genes across spatial locations.

**Figure 1:**
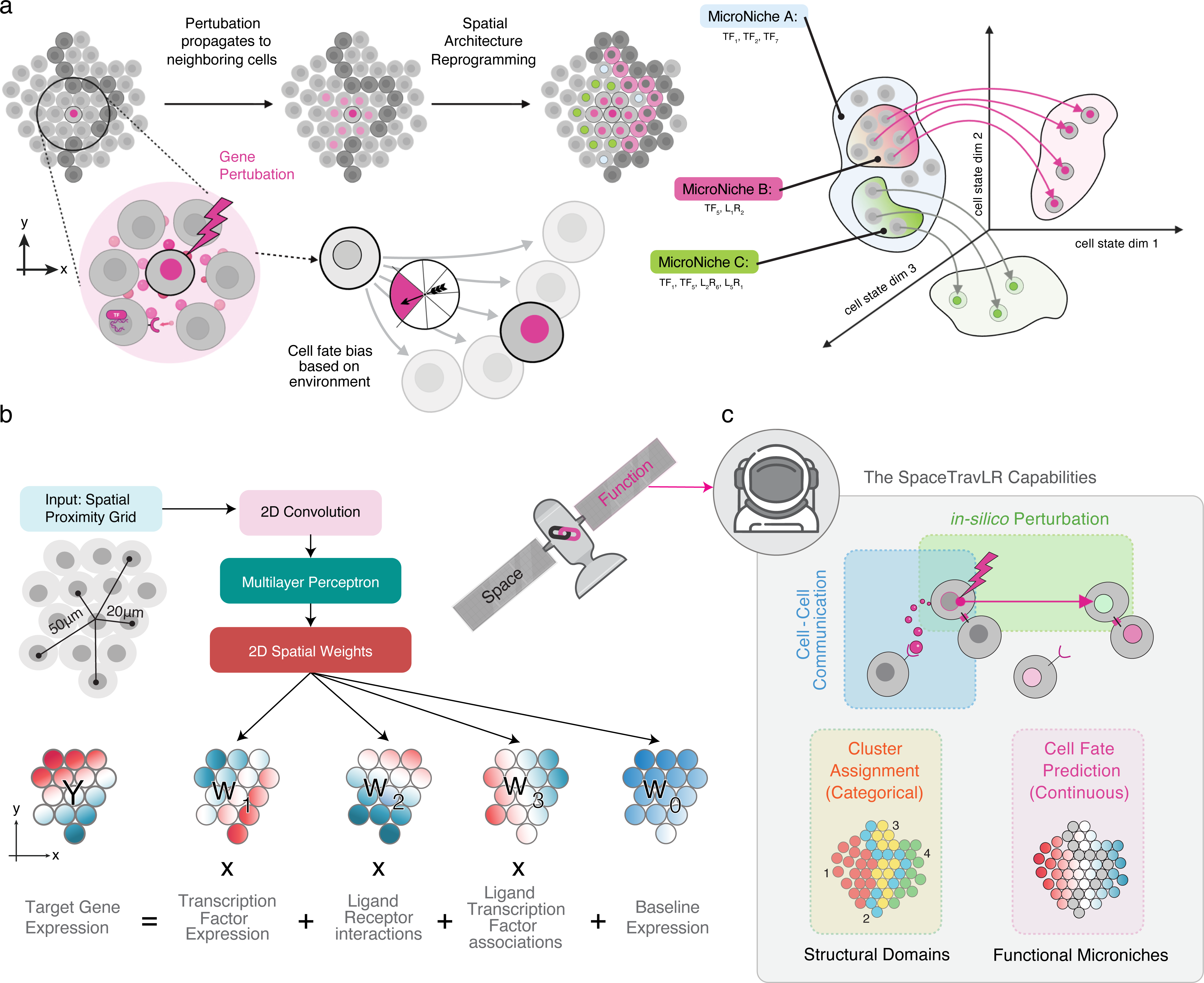
Linking spatial and functional biology with SpaceTravLR. **A.** Workflow of SpaceTravLR for finding functional microniches. Cell fate bias is dictated by both cell intrinsic and cell extrinsic signals. A perturbed cell can in turn perturb its neighbors via ligands. By delineating the tissue regions with distinct cell fates, we can identify functional microniches defined by their gene-gene interactions. **B.** SpaceTravLR performs 2D convolutions on the cell-cell spatial proximity input grid to generate spatially informed coefficients used in a regularized linear regression framework to predict the gene expression matrix. **C.** SpaceTravLR can perform traditional analysis on ST data such as *in-silico* perturbation, cell-cell communication inference and domain segmentation. By leveraging the spatial information and propagating perturbations from cell to cell, our model can then infer the functional consequences of the perturbations. Finally, we link space with function by uncovering regions with functionally different cell fate.

The model implements a scalable pipeline leveraging high performance computing for parallelized tensor computations to estimate the target gene expression based on spatially varying regulatory and signaling dynamics. Cell intrinsic regulation is captured through TF terms, while signaling is modeled via distance-weighted ligand expression from neighboring sender cells (optionally filtered by COMMOT^4^) to each receiver cell (**Fig. 1B**, Methods, **Supplementary Fig. 1**). Specifically, signaling is captured based on both ligand-receptor and ligand-TF associations. To integrate spatial information while maintaining biological interpretability, SpaceTravLR leverages convolutional neural networks^5^ (CNNs) to generate spatially resolved sparse graphs with differentiable edges. This architecture enables signals to propagate both within cells through regulatory edges and between cells through ligand–mediated connections, and is mathematically computed by efficient, gradient-based perturbation analysis via chain rule (**Supplementary Fig. 2**). We validate our approach on simulated data, accurately recovering the spatially varying coefficients for the three groups of modulators (**Supplementary Fig. 3**).

Once trained, SpaceTravLR is uniquely able to predict how cell intrinsic and cell extrinsic effects of perturbations vary spatially. We applied our model to multiple datasets across diverse spatial transcriptomic technologies and tissue types, showcasing wide applicability across different tasks such as (1) inferring functional cell-cell communication events, (2) *in-silico* modeling of functional and spatial reprogramming following changes in TFs, ligands and receptors expression, (3) identifying spatial domains and functional microniches and their defining genes (**Fig. 1C**). We applied SpaceTravLR to different biological systems to evaluate whether it can recapitulate known regulatory and signaling effects and uncover novel context-dependent mechanisms in tissue. Our entire pipeline (code and detailed documentation) and pretrained models across tissue types are made freely available as a resource for the community.

### SpaceTravLR captures the impact of individual and combinatorial perturbations on spatially resolved cell-intrinsic and cell-extrinsic circuits across tissue types

We first sought to elucidate whether SpaceTravLR can recover established regulatory and signaling effects using a spatial transcriptomic dataset from human tonsils generated using the Slide-tags^6^ platform. Tonsils have well-defined spatial architectures and include complex mixtures of immune cells (CD8 T cells, different B and CD4 T cell subsets, **Fig. 2A**) that work synergistically to mediate adaptive responses in a range of contexts spanning natural and vaccine-mediated immunity^7–9^. Thus, characterizing functional niches underlying these responses is of significant biological interest. We used this dataset to assess the model’s ability to capture how individual and combinatorial perturbations influence spatially resolved molecular networks across tissue types. Then, we extended this analysis to delineate spatial regions with distinct functional profiles, defined by differences in perturbation impact.

**Figure 2:**
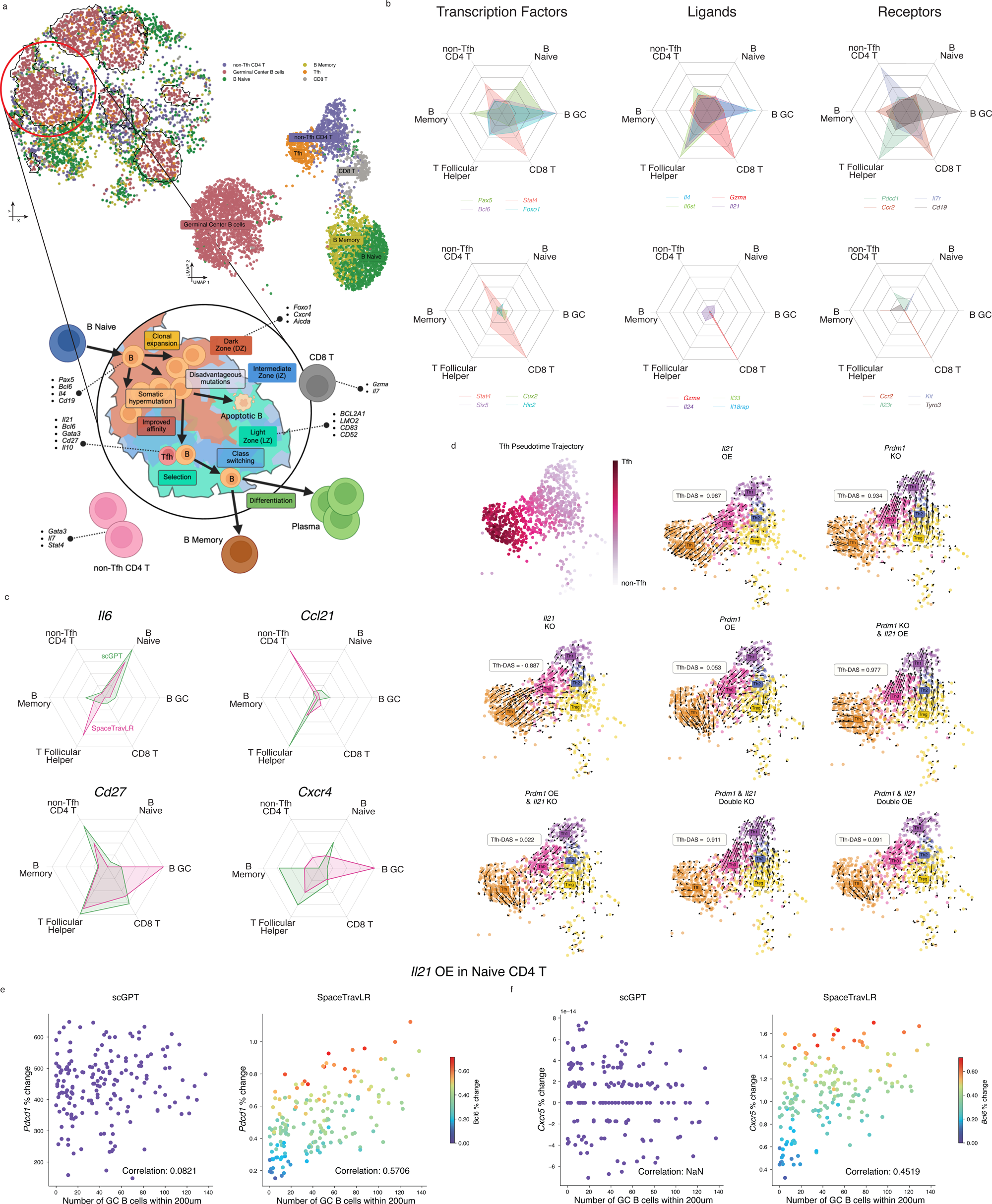
SpaceTravLR recapitulates the expected transcriptomic outcomes and spatial dependencies in diverse cell types within a complex tissue environment. **A.** Spatial organization and UMAP projection of the 6 major cell types in the Slide-tags tonsil dataset with distinct germinal center structures. Schematic illustrates the current biological understanding of key regulatory genes in these cell types and their interactions. **B.** Predicted relative magnitude of separate transcription factor, ligand, and receptor knockouts on each cell type (top panel). Genes not recognized as important to any of the 6 cell types are provided as a negative control, whose magnitude is scaled relative to the gene with the smallest impact among the displayed key genes (bottom panel). **C.** Relative magnitude of gene knockout effects on each cell type predicted by SpaceTravLR, CellOracle, and an adapted scGPT model for select ligands (top) and receptors (bottom). **D.** Tfh differentiation trajectory inferred using diffusion pseudotime, with higher values approaching 1 indicating more advanced stages of Tfh cell differentiation. SpaceTravLR simulated transition vectors showing the alignment of Tfh pseudotime trajectory with IL21 and PRDM1 KO and OE, and their combinations. **E-F.** Spatial dependencies of *Il21* OE on Naive CD4 T cells for scGPT (left) and SpaceTravLR (right) on E. *Pdcd1* and F. *Cxcr5* with respect to the number of GC B cells within 200um.

SpaceTravLR captured the spatially varying and cell-type dependent effects on cell-intrinsic and cell-extrinsic circuits across germinal center (GC) B cells, naive and memory B cells, plasma cells, CD8 T cells, and multiple CD4 T subsets. We quantitatively evaluated how to perturb a range of molecules across different cell types. The selected molecules included lineage-defining TFs and key ligands and receptors that are known to shape their cellular identities. In all cases, we accurately captured the cell-type-specific responses to relevant TFs, ligands and receptors with high sensitivity and specificity, demonstrating its ability to resolve both cell-intrinsic and signaling-driven perturbation effects (**Fig. 2B, top 3 panels**). For example, perturbation of the transcription factor *Pax5* most significantly impacts the three B-cell subtypes (B naive, B memory, and GC B, **Fig. 2B, top 3 panels**)10. We also accurately predicted that the ligands *Il4* and receptor *Pdcd1* knockouts (KOs) were most disruptive for Tfh cells, while the transcription factor *Bcl6* exerted a dual influence on both GC B cells and Tfh cells (**Fig. 2B, top 3 panels**)^11–22^. To assess the model’s specificity, we compared these findings with perturbations of TFs, ligands and receptors not expected to influence these cell fates and observed minimal effects (**Fig. 2B, bottom 3 panels**, red is the positive control relevant to one of the cell fates, the other colors correspond to molecules that are not relevant to any of the cell fates).

Next, we benchmarked SpaceTravLR against scGPT^23^ and CellOracle^24^, two recent state-of-the-art perturbation methods. We chose these two approaches as representative examples of conceptual paradigms widely used for *in-silico* perturbations. scGPT is a single-cell foundation model that generates informative cell-level embeddings that can be adapted into a linear model for zero-shot genetic perturbation. CellOracle, on the other hand, is an interpretable linear model leveraging underlying GRN architecture, but is limited to predicting TF perturbations only. Importantly, neither method can incorporate complex spatial organization and cellular interactions that SpaceTravLR can leverage. Across all benchmarks, SpaceTravLR performed significantly better than these approaches, capturing known effects of TF, ligand and receptor perturbations that one or both scGPT and CellOracle failed to predict (**Fig. 2C, Extended Fig. 1A**). SpaceTravLR also reproduced the pleiotropic effects of perturbations across multiple cell types. *Il6*, a crucial cytokine for GC development, promotes Tfh differentiation and GC B cell activation^25–27^. SpaceTravLR correctly predicted that the deletion of *Il6* most strongly affected Tfh and naive B cells, while scGPT was unable to recognize its importance for the Tfh cell type (**Fig. 2C**). Similarly, for *Ccl21*, a potent chemoattractant and costimulatory factor for CD4 T cells, only SpaceTravLR accurately captured the multi-faceted effects of this perturbation^28^. Conversely, for perturbations with more restricted effects, SpaceTravLR accurately resolved cell-type-specific responses, particularly those emerging from spatial cellular organization. Our model also accurately assessed the importance of receptors such as *Cd27* and *Cxcr4* in specific germinal center B-cell subsets (**Fig. 2C**). These receptors play critical roles in germinal center organization and B cell trafficking, and SpaceTravLR successfully recapitulated these structural relationships through its integration of spatial context. In contrast, scGPT failed to accurately capture the importance of key genes across distinct B-cell subsets^29–32^.

We also evaluated how well SpaceTravLR’s predictions function across different human and murine tissue types. SpaceTravLR accurately captured the impact of a wide range of TFs, ligands and receptors across murine kidney, human melanoma and murine lymph node samples (**Extended Fig. 1B-D**). Our method consistently captured the relative magnitude of gene knockout effects on the resident cell populations across all tissue contexts. These analyses also demonstrated SpaceTravLR’s robustness across different spatial transcriptomics platforms with varying resolutions from multi-cell to single cell or nuclei including XYZeqV2^33,34^, Slide-tags^6^, and Slide-seqV2^35^. Despite variation across platforms, SpaceTravLR performed effectively on both single-nuclei data and lower-resolution spots containing mixed transcriptional signals from multiple cells.

Next, we sought to evaluate SpaceTravLR’s ability to assess the quantitative effects of combinatorial gene perturbations, within a complex regulatory and signaling network. Because of our model’s unique formulation, we can simulate previously untested perturbations for any ligand-receptor or ligand-transcription factor pairs. As a proof of concept, we modeled perturbation combination of *Prdm1* and *Il21* with opposing roles in Tfh differentiation. *Prdm1* is a transcriptional repressor of the Tfh lineage, while *Il21* is a cytokine that promotes it^36–40^. By testing both perturbations separately or together, we can rigorously assess the model’s capacity to capture how activators or repressors work within shared regulatory and signaling circuits. We quantitatively assessed our predictions by aligning the inferred perturbations along diffusion pseudotime (where the Tfh-differentiation alignment score (Tfh-DAS) (−1 to +1) represents the extent to which a perturbation inhibits or promotes progression along the Tfh trajectory) (**Extended Fig. 2A**). First, consistent with previous studies, SpaceTravLR predicted the expected effects: *Il21* over-expression (OE) and *Prdm1* KO promoted Tfh differentiation as indicated by high Tfh-DAS, while the opposite perturbations (*Il21* KO and *Prdm1* OE) suppressed it, giving low Tfh-DAS (**Fig. 2D**). Interestingly, *Prdm1* deletion also induced a partial Th1 fate, consistent with the existence of a shared Th1/Tfh developmental trajectory^41–43^ (**Fig. 2D**). Next, we looked at OE and KO combinations of *Il21* and *Prdm1*, including those that are antagonistic. The synergistic combinations of *Prdm1* KO and *Il21* OE had the greatest impact on promoting the Tfh lineage while *Prdm1* OE and *Il21* KO inhibited the Tfh lineage (**Fig. 2D**). But more interestingly, of the antagonistic combinations, *Prdm1* had a dominant role. *Prdm1* and *Il21* KO promoted the Tfh lineage while *Prdm1* and *Il21* OE repressed the Tfh lineage (**Fig. 2D)**. This agrees well with the known signaling circuit of *Prdm1* (a TF) being downstream of *Il21* (a cytokine)^40^. Intriguingly, the antagonistic combinations had less dramatic impacts on cell fate than synergistic combinations, demonstrating that our method accurately quantitatively identifies these cell fates (**Fig. 2D**).

^40^.Next, we moved beyond the impact of individual and combinatorial perturbations to examine how the spatial context influences cell fate, a critical step in defining spatial functional microniches in tissues. Tfh cells need continual extrinsic signals from GC B cells mediated via Cd40/Cd40l interactions^37,44^. We examined the spatially-resolved combinatorial impact of three molecules promoting the Tfh lineage – *Bcl6* (a TF), *Pdcd1* (a receptor) and *Il21* (a ligand) Interestingly, when we modeled*Il21* OE in naive CD4 T cells, *Bcl6* and *Pdcd1* expression positively correlated with the number of GC B cells within a 200 micron spatial neighborhood (**Fig. 2E**). This demonstrates the existence of functional microniches where *Bcl6*, *Il21,* and *Pdcd1* synergistically promote Tfh fate only when sufficient GC B cells are present in proximity. Similar spatial dependencies were also observed for *Bcl6*, *Il21*, and Tfh lineage promoting receptor, *Cxcr5*^45^ (**Fig. 2F**). scGPT completely failed to identify this critically important biological dependency (**Figs. 2E, 2F**), and CellOracle is fundamentally incapable of modeling this as it can only perturb TFs. Thus, SpaceTravLR is the first method capable of individual and combinatorial gene perturbations to resolve spatially dependent regulatory and signaling (cell-cell communication) effects.

### Dissecting cell fate decisions dictated by the local environment

While existing *in-silico* methods can predict the differential impact of genetic perturbations across cell types/states, these methods are unable to uncover alternate cell fates for the same cell type/state based on spatial location^46^. We used SpaceTravLR to infer the impact of genetic perturbations in modulating cell states across regions of varying spatial organization in complex tissues. This helps define functional microniches as changing spatial environments can reshape cell type and behavior.

We focused on GC B cells, which cycle between dark zone (DZ) and light zone (LZ) through an intermediate zone (iZ) as they undergo affinity maturation within secondary lymphoid tissues^9,47^. In the DZ, B cells proliferate and undergo somatic hypermutation (SHM), to increase antibody diversity and affinity, then migrate to the LZ to capture antigens to present to Tfh cells. High-affinity B cells receive survival signals and differentiate, while low-affinity clones return to the DZ for additional rounds of mutation and selection^48^. A key TF driving this process in both GC B cells and Tfh cells is *Foxo1* (**Fig. 3A**). We sought to explore whether SpaceTravLR could predict distinct spatial outcomes of *Foxo1* perturbations within GC compartments (DZ, LZ and iZ). SpaceTravLR predicted that *Foxo1* KO caused the DZ GC B cells to take on a LZ identity, but LZ GC B did not change their identity despite exhibiting a similar transcriptional profile to DZ GC B cells (**Figs. 3B, 3C, Extended Fig. 2B)**. This result aligns closely with the functional role of *Foxo1* characterized in murine models which demonstrate experimentally that *Foxo1* is critical for the spatial polarization and structure of the GC, enabling GC B cells to re-enter the DZ, and that its loss diverts the cells to the LZ^14,49,50^. This demonstrates the ability of SpaceTravLR to accurately predict cell state changes due to altered spatial migration dynamics (**Figs. 3B, 3C**). Furthermore, the degree to which a cell is impacted, as quantified by its transition probability, is determined primarily by its spatial location (**Extended Fig. 2B, Supplementary Fig. 4**). SpaceTravLR correctly inferred that DZ GC B cells with similar transcriptomes can have different transition probabilities due to their spatial niches.

**Figure 3:**
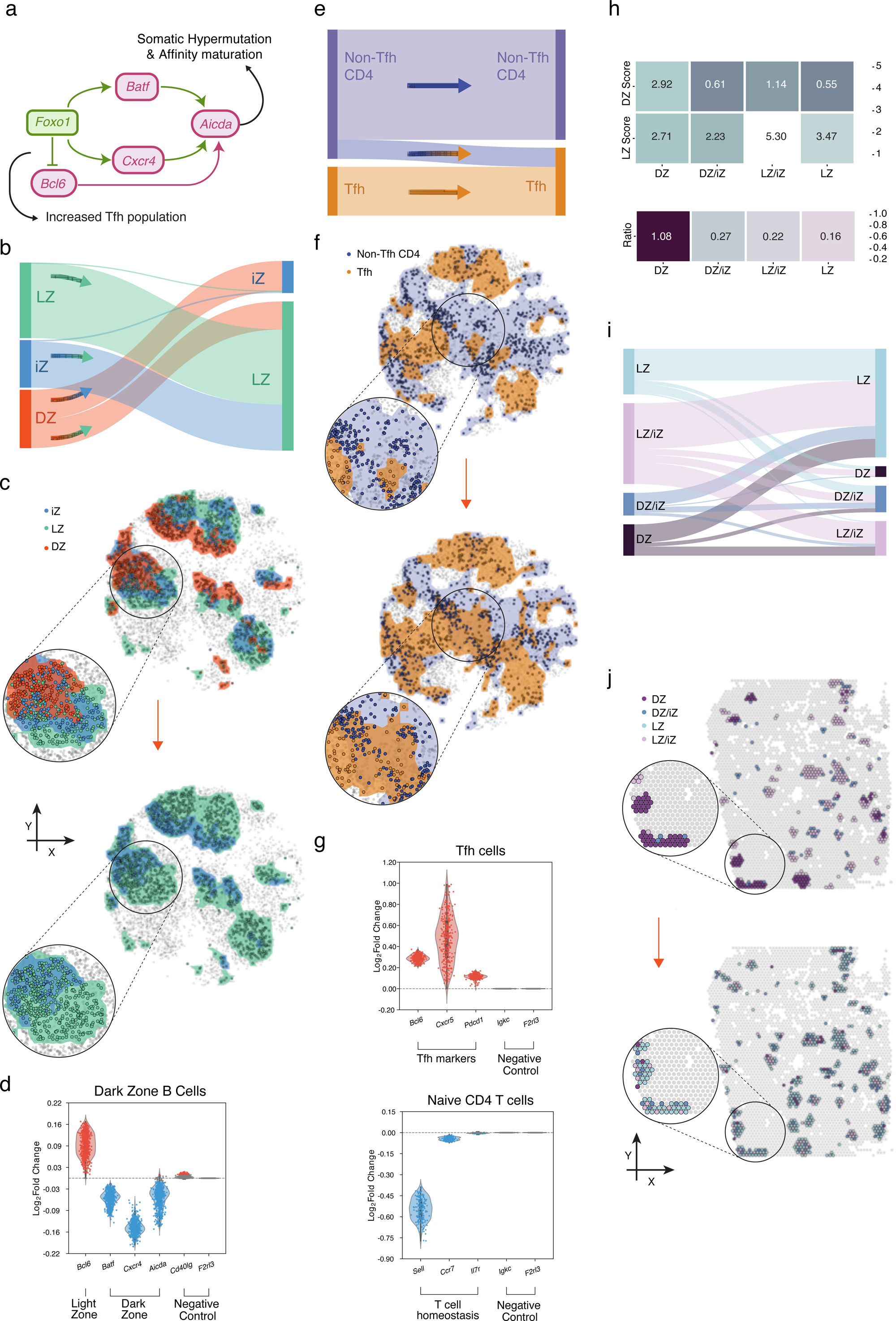
SpaceTravLR captures spatial cell-to-cell variability in perturbation response within germinal centers. **A.** Simplified subnetwork of *Foxo1* previously established in the litterature. *Foxo1* positively regulates the expression of *Batf* and *Cxcr4* which are essential for the expression of *Aicda*, an essential gene for somatic hypermutation in GC dark zones. *Foxo1* has also been linked to the survival of naive CD4 T cells while its deletion has been confirmed to amplify Tfh populations. **B.** Representation of the proportion of transitions between GC B cells in the dark zone, marginal zone, and light zone compartments resulting from *Foxo1* KO. **C.** Spatial visualization of the impact of *Foxo1* KO on the GC B cells. **D.** Simulated log2 fold change of key GC genes following *Foxo1* KO **E.** Quantification of CD4 T cell transitions between Tfh and non-Tfh states resulting from *Foxo1* KO. **F.** Spatial visualization of the impact of *Foxo1* KO on the Tfh and non-Tfh cells **G.** Simulated log2 fold change of relevant genes downstream of *Foxo1* per cell in Tfh (top) and Naïve CD4 T cells (bottom) **H.** Cell2location scores for GC light zone and dark zone B cells (top) and their proportions (bottom) across the 4 Leiden clusters representing the most GC-like Visium spots. **I.** Representation of the proportion of transitions between the 4 Leiden clusters from *Foxo1* knockout. **J.** Spatial visualization of the impact of *Foxo1* KO on the composition of GC spots in the test sample Visium lymph node.

Critically, the predicted effects of *Foxo1* KO on tissue architecture were both interpretable and biologically meaningful. They were driven by alterations in other key genes and regulators that are known to control GC dynamics. *Foxo1* positively regulates *Batf* and *Cxcr4*, which are essential for somatic hypermutation in the DZ through *Aicda*^14,49^ (**Fig. 3A**). SpaceTravLR correctly simulated the increased expression of *Bcl6*, and decreased expression of *Batf* and *Cxcr4* in *Foxo1* KO (**Fig. 3D**), demonstrating gene-level precision and robust network propagation. SpaceTravLR achieves this level of mechanistic accuracy (established in murine models) with no explicit encoding of biological priors but solely based on interpretable model architecture which leverages the underlying biological network. These phenotypic and mechanistic changes identified by SpaceTravLR were not captured by CellOracle (**Extended Figs. 2C-F**), which failed to identify known downstream genes in the *Foxo1* circuit (**Extended Fig. 2F**). SpaceTravLR’s incorporation of signaling, particularly *Cxcr4*, proved critical for capturing the impact of *Foxo1* knockout on *Aicda* expression^32^ (**Extended Figs. 2G, 2H**).

Though *Foxo1* is best known for its role in B cells, its regulatory effects on CD4 T cells are also well characterized. In mice, *Foxo1* knockout is known to remove repression of *Bcl6*, resulting in an expanded population of *Cxcr5*+ *Pdcd1*+ Tfh (bonafide GC Tfh) cells^44,51,52^. Again, SpaceTravLR can predict just using the Slide-tags data, that upon *Foxo1* KO, the population of Tfh cells increases, as non-Tfh take on a Tfh transcriptional identity, but localization is largely unchanged (**Figs. 3E, 3F).** This occurs because of a concomitant increase in the expression of *Pdcd1*, *Cxcr5*, and *Bcl6* (**Fig. 3G**). In naive CD4 T cells, *Foxo1* has been linked to homing and survival of naive T cells by regulating *Sell*, *Ccr7* and *Il7r*^53^. Simulated *Foxo1* knockout causes the downregulation of all three genes, consistent with experimental evidence (**Fig. 3G**). Additionally, CD4 T cells activated in the presence of TGF-β typically differentiate into *Foxp3*+ regulatory T cells, but *Foxo1* knockout redirects this differentiation toward *Ifng+* Th1 cells instead^52,54^. Our *in-silico Foxo1* perturbation captured this phenotypic switch, demonstrating decreased expression of Treg-associated genes (*Ccr4*, *Foxp3*, **Extended Fig. 2I**), while showing increased expression of Th1-associated genes (*Ifng*, *Cxcr3*, **Extended Fig. 2I**).

These results utilized a CNN-based vision model which explicitly learns spatially dependent gene-gene relationships but is distinct from a transformer which may enhance learning spatially dependent relationships. Given the structural complexity of the GC, we sought to evaluate whether replacing our CNN-based vision model with a more sophisticated Vision Transformer (ViT) would enhance performance. Both architectures learned highly similar gene-gene relationships and achieved comparable performance in predicting target gene expression (**Supplementary Fig. 5**). Given their equivalent predictive accuracy, we selected the CNN-based model due to its substantially lower parameter count (i.e., less prone to overfitting even on sparse data increasing generalizability of our model across datasets) and reduced training costs. Importantly, our CNN-based architecture effectively captures and incorporates spatial information at the level of other state-of-the-art vision models. To minimize spurious associations within our linear model, we employ group-lasso regularization (**Supplementary Figs. 6, 7**). Finally, we selected the depth of network propagation based on overall transcriptomic convergence and established findings regarding network topology (**Supplementary Figs. 8, 9**). Thus, we find CNN is sufficient to capture spatially dependent relationships in our model.

Overall, the regulatory and signaling networks learned by SpaceTravLR generalize across different tissues and spatial profiling technologies. Using the relationships learned in the tonsil tissue, our model accurately predicts the effects of gene perturbations in an unseen lymph node sample. Furthermore, while the tonsil was profiled with Slide-tags, the lymph node was profiled with Visium^55,56^, a platform that captures a mixture of cells in one spot. Regardless, SpaceTravLR is still able to predict that *Foxo1* knockout in the lymph causes a decrease in the spots with higher proportions of DZ cells, consistent with the shrinking of dark zones in experimental observations (**Figs. 3H-J**). The cross-context (across related biological tissues – tonsil and lymph node) and cross-platform generalizability of SpaceTravLR make it broadly applicable to capturing cell-intrinsic effects that lead to spatially resolved functional microniches.

### Simulating functional paracrine signaling at single-cell resolution

While cell-intrinsic effects described above are a critical facet of signaling systems, they are often not sufficient to fully explain distal effects in tissue. SpaceTravLR also enables modeling of cell-extrinsic paracrine signaling networks, a critical and often underexplored facet of tissue organization. Existing *in-silico* methods for investigating cell-cell communication and microniche discovery often operate independently, overlooking a key biological principle that microniches shape cell function and behavior through intercellular signaling. To address this, we evaluated whether SpaceTravLR could capture paracrine signaling by isolating a perturbation in one cell type and quantifying its downstream impact on a spatially adjacent, unperturbed population. This is a complex problem of major biological significance not currently addressed by existing *in-silico* perturbation methods or foundation models.

For this, the kidney offers a great system for studying functional microniches and immune–microenvironment interdependence since the medulla contains varying regions of osmotic gradients. These hypertonic niches drive epithelial cells to produce chemokines that recruit resident and monocyte-derived phagocytes, and bias macrophages toward pro-inflammatory phenotypes under salt challenge. Here we set out to study how epithelial-immune cell interactions define and maintain different microniches. A functional understanding of cell-cell communication requires modeling the causal impact of ligand–receptor interactions on cell state at a single cell level. In collaboration with Survey Genomics, we developed a true single cell spatial profiling assay, XYZeqV2, based on a spatially hashed nanowell technology. Briefly, by spatially barcoding a fresh slice of tissue prior to dissociation, we are able to recover the position of each cell using standard single cell workflows such as 10x. Thus, we can take advantage of the spatially resolved single cell data to probe cell-type specific gene expression and spatially contextualize receptor–ligand interactions. The functional relevance of the hypertonic macrophages identified here is their expression of inflammasome sensors such as Nlrp1b which responds to high-salt conditions and subsequently triggers IL-1β release during cellular stress^56,57^ (**Extended Fig. 3A**). These hypertonic macrophages also act as salt sensors under homeostatic conditions, and alterations in their functions are directly related to delayed wound healing, and aberrant responses in infection and chronic disease such as atherosclerosis^57^.

Macrophage migration inhibitory factor *Mif* is a key proinflammatory cytokine and chemokine-like function cytokine that is secreted predominantly from epithelial cells upon inflammatory and stress stimulation (**Fig. 4A**). Spatial mapping confirmed that these hypertonic macrophages are localized predominantly in the central kidney regions, where osmotic gradients are steepest, while regular macrophages occupy the periphery (**Fig. 4B**). These distinct populations appeared in other murine kidney replicates (**Supplementary Fig. 10**). As such, we predicted that the influence of *Mif* would be strongest within the central regions of the kidney, where the hypertonic zones were most apparent. In our murine kidney sample, *Mif* is primarily expressed by epithelial cells acting through several cell surface receptors expressed by other cell types, most notably myeloid cells, while *Cd74*, a high affinity receptor for *Mif*, is predominantly expressed by macrophages (**Fig. 4C**). Using SpaceTravLR, we modeled the impact of *Mif* KO in epithelial cells. SpaceTravLR successfully captures the signaling between *Mif* and *Cd74* and predicts substantial transcriptional changes in the macrophage population upon *Mif* KO only in the epithelial cells (**Fig. 4D**). Consistent with our hypothesis, the *in-silico* knockout caused the hypertonic macrophages to adopt transcriptomic profiles resembling regular macrophages. Furthermore, not all hypertonic macrophages were equally affected. Cells located in the central regions of the kidney tissue were most likely to transition towards regular macrophages following *Mif* KO, even when they had distinct transcriptional states (**Extended Fig. 3B**). This illustrates the ability of SpaceTravLR to capture spatially dependent cell-extrinsic signaling circuits.

**Figure 4:**
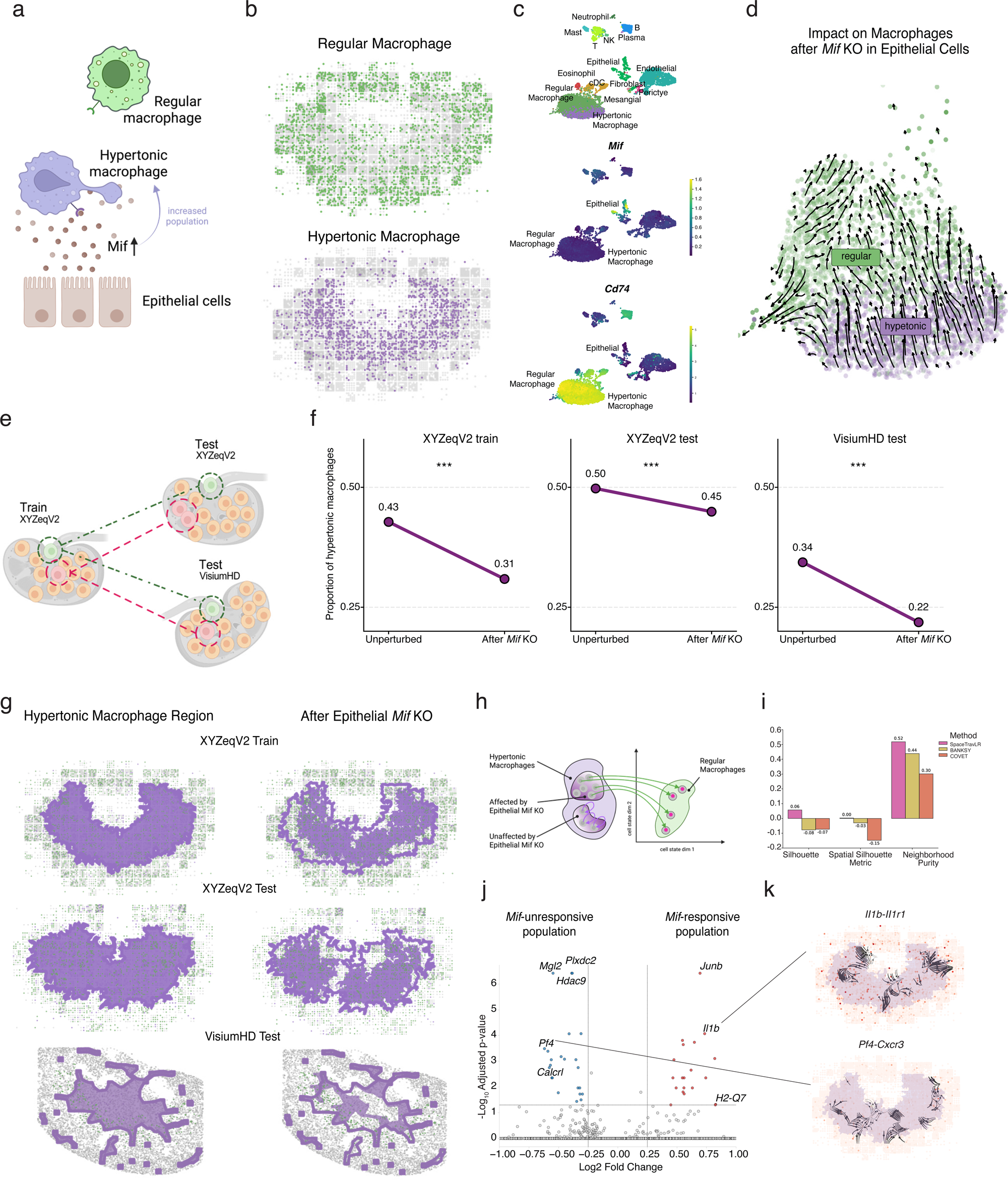
SpaceTravLR infers the functional consequences of cell-cell communication events at true single cell resolution in murine kidney. **A.** Schematic of paracrine signaling between epithelial cells and macrophages. Epithelial cells secrete Mif, which promotes the hypertonic macrophage population. **B.** Hypertonic macrophages localize to the inner regions of the kidney, while regular macrophages mostly populate the outer regions. **C.** UMAP of kidney single-cell data colored by cell type (top), *Mif* expression intensity (middle), and *Cd74*, one of the receptors for *Mif* (bottom). **D.** Vector plot showing the simulated transitions between the hypertonic and regular macrophage populations in UMAP space. **E.** Cross-platform validation approach. The model trained on XYZeqV2 data is generalized to another XYZeqV2 sample and a publicly available VisiumHD sample. **F-G.** Quantification and spatial effects of Mif knockout in epithelial cells on the hypertonic macrophage population proportion. **H.** Schematic of paracrine signaling-driven cell fate alterations in hypertonic macrophages, identifying two distinct groups: *Mif*-responsive and *Mif*-unresponsive populations. **I.** Validation of SpaceTravLR’s partitioning against state-of-the-art microniche methods (BANSKY and COVET) using silhouette score, spatial silhouette score, and spatial neighbor purity metrics. **J.** DEGs between SpaceTravLR’s *Mif*-responsive and *Mif*-unresponsive partitions, including several inflammatory gene markers. **K.** COMMOT Cell-cell communication analysis for *Pf4* and *Il1b*, two DEGs from this comparison.

After training on a single kidney sample, our model can be applied to unseen samples. We used the Spatial-Linked Alignment Tool (SLAT)^58^, a graph-based algorithm that enables the mapping of cells across heterogeneous slices by incorporating both spatial location within the tissue and gene expression profiles. We apply SpaceTravLR model weights from the training sample to two new unseen samples to generate predictions (**Fig. 4E**, **Methods**). The new samples include one profiled using the XYZeqV2 platform, and the other using VisiumHD. In both cases, simulated epithelial-cell *Mif* knockout induced a portion of hypertonic macrophages to adopt regular macrophage identities. Despite differences in spatial resolution and cell-type composition across these datasets, SpaceTravLR consistently predicted that *Mif* depletion reshapes the spatial organization of macrophage populations, demonstrating its ability to generalize across technologies and samples (**Figs. 4F, 4G**).

Beyond capturing spatially dependent paracrine signaling, SpaceTravLR also identified two functionally distinct populations within hypertonic macrophages - *Mif*-responsive and *Mif*-unresponsive (**Fig. 4H**). We evaluated whether BANSKY^59^ and COVET^60^, two state-of-the-art structural niche identification methods were also able to identify these populations. For BANSKY, we examined only the clusters (6 out of 16) that contained hypertonic macrophages (**Extended Fig. 3C**). For COVET, we computed embedding using all cells, then applied Phenograph^61^ clustering to just the hypertonic macrophages (14 clusters, **Extended Fig. 3D**). Unlike SpaceTravLR, neither method captures which hypertonic macrophages are responsive or unresponsive to *Mif*. Furthermore, beyond being biologically interpretable, the subsets identified by SpaceTravLR outperform both methods in three complementary clustering metrics: silhouette score, which measures how well-separated clusters are in gene expression space; spatial silhouette score, which evaluates cluster separation in physical space using spatial coordinates; and neighborhood purity, which quantifies local spatial coherence by measuring the fraction of each cell’s spatial neighbors that share the same cluster assignment. SpaceTravLR outperformed BANSKY and COVET across all spatial and transcriptomic evaluation benchmarks but also produced spatial clusters that were more biologically interpretable (**Fig. 4I**). This interpretability was further supported by differentially expressed gene (DEG) analysis, which revealed numerous inflammation-related genes with distinct functional roles between the two SpaceTravLR-defined partitions (**Fig. 4J**). *Il1b* was upregulated in the *Mif-*responsive population, demonstrating that within hypertonic macrophages, the degree of *Il1b* expression is indicative of the level of *Mif* responsiveness. Conversely, *Pf4* showed higher expression in *Mif*-unresponsive hypertonic macrophages. Blockage of the *Pf4* signaling pathway has been linked to reduced inflammation, suggesting a role in an inflammatory macrophage phenotype^62–64^. These two DEGs between the two groups of hypertonic macrophages have complementary effects through likely separate inflammatory circuits through cell-cell communication and highlight the model’s ability to identify functionally relevant partitions specifically related to *Mif* signaling (**Fig. 4K**). Together, these results suggest that SpaceTravLR captures potential downstream consequences of *Mif* perturbation and can predict how local signaling shapes functionally distinct macrophage subsets.

We next asked whether our model could generalize this framework to paracrine signaling involving a more promiscuous ligand. In our murine kidney sample, *Tgfb2* is highly expressed by endothelial, epithelial, and mesenchymal stromal cells, while *Tgfbr1*, a component of the corresponding Tgfβ receptor complex, is most highly expressed in myeloid cells (**Extended Fig. 3E**). We performed *in-silico Tgfb2* knockouts in each of the three source cell types that express the ligand to measure the transcriptional outcomes in myeloid cells. The predicted effects varied depending on the perturbed cells (**Extended Fig. 3F**). Importantly, the extent of the perturbation is not solely dependent on the magnitude of expression of ligand or receptor in any given cell type, but rather on overall spatially dependent signaling across cell types. For example, upon *Tgfb2* KO there were significant differences in epithelial-mesenchymal stromal interactions despite *Tgfbr1* and *Tfgb2* not being expressed at very high levels in these cell types. Overall, our analyses illustrated the ability of SpaceTravLR to capture complex spatially dependent signaling and quantify the variable impact, revealing how the same genetic perturbations can produce divergent paracrine signaling across neighboring cell types and states.

### Modeling emergent cell fates and spatial perturbation outcomes

Next, we evaluated whether SpaceTravLR could model cellular states beyond those represented in the profiled transcriptome which enables us to understand emergent cellular behaviors not captured in the data. We first applied the model to a Slide-seqV2 mouse embryo dataset^65^, where *Tbx6-*deficient embryos classically develop ectopic neural tubes instead of somatic cells (**Figs. 5A, 5B).** *Tbx6* is a transcription factor expressed in the presomitic mesoderm (PSM) and is required for somite partitioning and specification. To test whether our model could recover this developmental phenotype from spatial transcriptomics data, we trained it on published WT embryo data and then performed *in-silico Tbx6* KO across all the cells at embryonic day 9.5 (E9.5), a well-studied stage critical to neural and brain development. Our model accurately predicted that *Tbx6* depletion would most significantly alter^66^ the cell fate of somites, neuromesoderm progenitors (NMP), and PSM in embryos (**Figs. 5C, 5D**). Overlaying the ectopic tube transcriptomic profile, extracted from the *Tbx6* KO Slide-seqV2 dataset, onto the simulated perturbation transcriptomic profile (**Figs. 5C, 5D**), we observe this shift towards the ectopic phenotype. Our results demonstrate that SpaceTravLR can accurately infer the emergence of previously unseen cell states directly from ST data.

**Figure 5:**
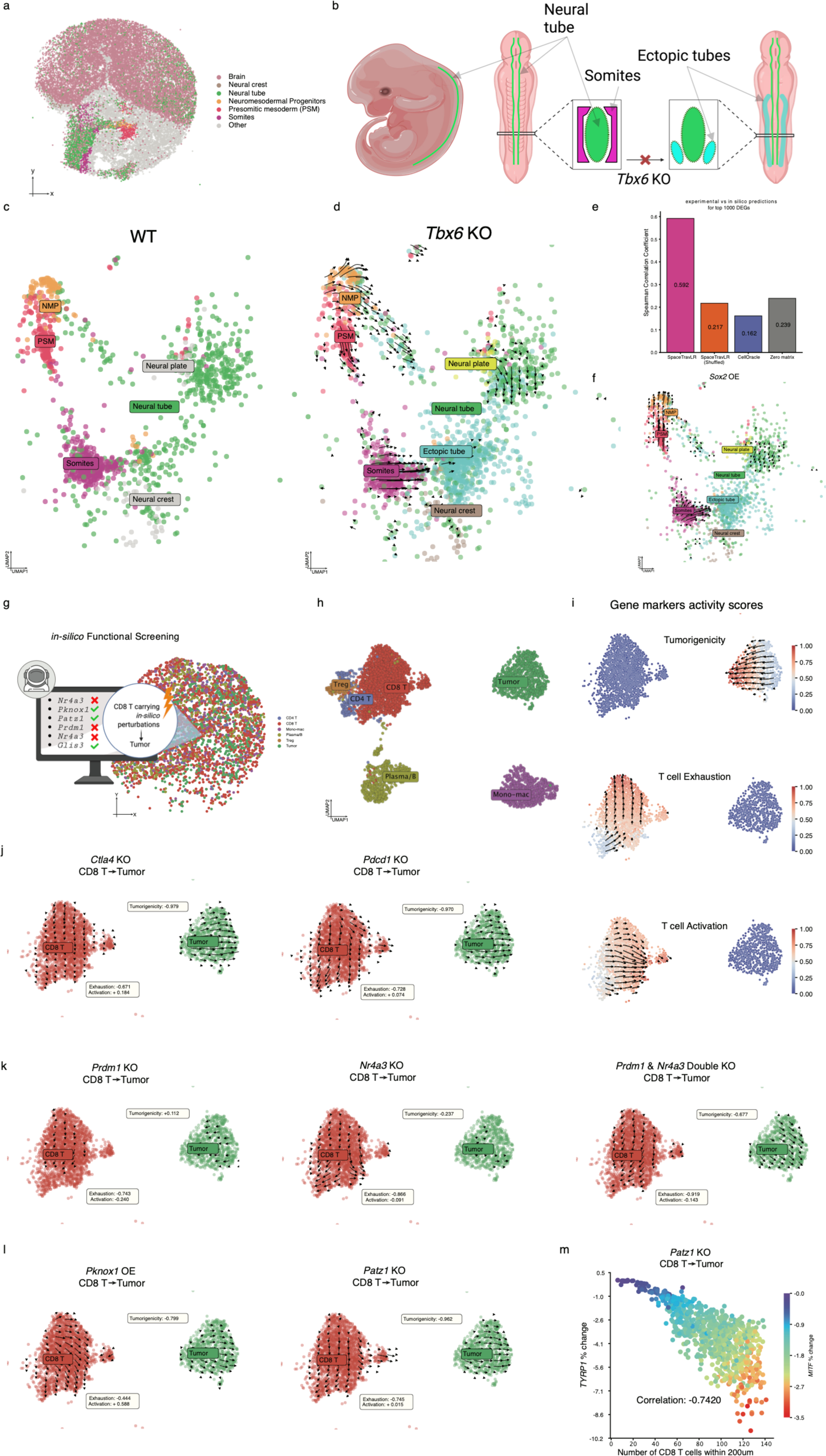
SpaceTravLR enables discovery in highly dynamic and disorganized tissues. **A.** Spatial map of E9.5 WT Slide-seqV2 data and different cell types used for training. **B.** Schematic showing the expected somitic and neural developmental phenotype in WT embryo and its alterations after *Tbx6* KO. **C.** UMAP of cell types in during normal development. **D.** Ectopic tube cells from the real *Tbx6* knockout sample are projected into the same UMAP space as the wildtype cells. Vectors demonstrate SpaceTravLR predicted cell-state transitions resulting from *Tbx6* knockout. **E.** Schematic for *in-silico* functional screening in Slide-tags melanoma sample. **F.** UMAP showing annotated cell types in the melanoma sample. **G.** Gene signature scores overlaid on UMAP projection to show trends across the clusters. **H.** UMAP projection of cell types in the melanoma sample. **I.** UMAP projection of only the CD8 T cells and tumor cells colored by gene marker scores for tumorigenicity (*Pmel, Mitf, Dct, Cdk2, Cdh1, Tyrp1, S100b*), T cell exhaustion (*Tigit, Lag3, Tox, Btla, Havcr2*), and T cell activation (*Itga1, Gzma, Gzmk, Ccl5, Ccr5*). Gene marker activity vectors indicate the direction of the steepest increase for each score, pointing toward cell states with the highest scores. **J.** Impact of CD8 T cell-specific *Ctla4* and *Pdcd1* knockout on CD8 T cell and tumor cell states. Scores are computed from their alignment with the gene markers’ activity vectors. **K.** Individual *Prdm1* and *Nr4a3* knockouts have a less negative effect on tumorigenicity and exhaustion, and a more positive effect on activation compared to their combinatorial knockout. **L.** Alterations to novel targets, *Pknox1* OE and Patz1 KO, are predicted to have highly negative tumorigenicity and exhaustion scores and positive CD8 T cell activation scores. **M.** Gene expression changes in tumor cells following CD8 T cell-specific *Patz1* knockout. Tumor cells with more CD8 T cells within 200 μm show significantly greater downregulation of *Tyrp1* and *Mitf*. Tyrp1 percent change is strongly negatively correlated with local CD8 T cell density (r = −0.742).

Because *Tbx6* and *Sox2* form an antagonistic feedback loop^67^ that controls mesoderm to neural fate decisions, we simulated *Sox2* OE to test whether our model could infer this reciprocal developmental pathway. SpaceTravLR accurately predicted that *Sox2* OE drives somites to reacquire progenitor-like features (**Figs. 5C, 5E and 5F**) and reproduced the expected shift in canonical neural tube patterning markers (**Supplementary Fig. 11**). Crucially, although the published KO data was generated at the zygote stage, our *in-silico* KO was performed at E9.5. Thus, while *Tbx6* KO more directly mirrors the observed phenotype, *Sox2* OE may be more faithful to the underlying developmental process, where the PSM fails to develop correctly and potentially becomes more NMP-like instead (**Fig. 5F**). Across the top 1000 DEGs between the WT and KO, SpaceTravLR significantly outperformed in magnitude and direction of the expression changes compared to existing state-of-the art perturbation models and other benchmarks (**Fig. 5E**).

To further test SpaceTravLR’s ability to infer regulatory and signaling relationships in complex and heterogeneous tissue, we applied our framework to human melanoma to systematically screen for perturbations that could alter immune cell-tumor interactions. Despite showing remarkable promise, checkpoint blockade immunotherapy is only durable for a small subset of patients. Several factors limit the success of T cell immunotherapy in solid cancers, including cell trafficking to the tumor, exhaustion due to chronic antigen exposure, and other immunosuppressive mechanisms operating within the tumor microenvironment (TME). Here, we sought to identify key genes in CD8 T cells, that restrain CD8 T cell exhaustion and promote anti-tumor phenotypes, by systematically perturbing all expressed TFs, ligands, and receptors. Using a Slide-tags human melanoma sample with a complex mixture of immune (monocytes, B, Treg, CD4 T, CD8 T) and tumor cells (**Fig. 5G**) we aimed to identify genes that could enhance CD8 T cell cytotoxicity while reducing tumorigenicity. We mapped marker genes to spatial vector fields to define three key metrics – tumorigenicity, T cell exhaustion, and T cell activation (**Figs. 5H, 5I, Methods**). In the tumor compartment, vectors aligned along the tumorigenicity axis capture transitions from less aggressive to more aggressive and invasive phenotypes. In the CD8 T cell compartment, analogous gradients reflect activation-to-exhaustion dynamics, with vector directions representing the continuum from naïve or effector-like to terminally exhausted states (**Figs. 5H, 5I**). We superimposed these cellular fate outcome vectors with the SpaceTravLR simulated spatially resolved perturbation vectors to find gene targets in CD8 T cells that simultaneously rescue exhausted T cells, enhance T cell cytotoxic activity and decreases tumorigenicity. This demonstrates SpaceTravLR’s utility as a screening tool for the identification of novel anti-tumor immunotherapy targets.

As expected^68,69^, KO of known checkpoint inhibitory receptors (*Ctla4* and *Pdcd1*) enhanced anti-tumor activity (**Fig. 5J**), confirming SpaceTravLR’s predictive accuracy. This is in line with their well-studied roles as *Ctla4* and *Pdcd1* and primary targets of current checkpoint inhibitors widely used in the clinic. We also looked at more complex combinatorial effects of perturbing TFs and receptors. Consistent with previously published experimental results^70^ in lymphoma, *Prdm1* KO reversed exhaustion, but failed to enhance T-cell cytotoxicity against tumors, likely due to the reported compensatory upregulation of *Nr4a3* (**Fig. 5K**). Also, consistent with our expectation, our simulation revealed that *Nr4a3 KO* marginally increased anti-tumor activity. However, consistent with expectation, *Prdm1* and *Nr4a3* double KO synergistically prevented T-cell exhaustion while improving anti-tumor outcome (**Fig. 5K**). Beyond known regulators, our model discovered putative novel targets, *Patz1* and *Pknox1,* that upon knockout, would confer high anti tumorigenicity, high activation and restrain exhaustion (**Fig. 5L**). Intriguingly, these effects were spatially restricted and the predicted magnitude of cytotoxic activity on the tumors, as measured by *Tyrp1* expression on the tumors, scaled in proportion with localized CD8 T cell density (**Fig. 5M, Supplementary Fig. 12**). Together, these results demonstrate SpaceTravLR’s ability to generalize across developmental and disease contexts to model regulatory and cell intrinsic and extrinsic signaling networks in the tissue space. More importantly, the model permits unbiased virtual functional screens that are quick, scalable, accurate and a tiny fraction of the cost of performing them experimentally to uncover actionable perturbations that can potentially alter cell fate and transitions within the multicellular spatial neighborhoods.

### SpaceTravLR discovers a novel Ccr4-dependent spatially restricted Th2 microniche that influences lung migration in allergic asthma

Finally, we aimed to use SpaceTravLR to understand spatial cell dynamics in a setting where we could experimentally validate results. We focused on a murine model of allergic asthma (**Fig. 6A**) to elucidate the functional roles of key molecules in determining spatial migration patterns of T cell subsets. Allergic asthma is driven by a population of pathogenic Th2 cells that migrate from the mediastinal lymph nodes (medLNs) to the lung^71^. Specifically, our prior work demonstrates that Th2 cells require signaling mediated by synergy between *Il10* and *Il2* and its corresponding receptors – *Il2ra, Il2rb,* and *Il2rg*^72^. Here we applied SpaceTravLR to recapitulate this mechanism de-novo as well as find novel molecules underlying the spatial migration of these pathogenic cells (**Figs. 6A, 6B**). We specifically analyzed a Slide-seqV2 dataset with paired TCR-seq collected from the medLN three days after adoptive transfer of HDM-specific TCR transgenic CD4 T cells followed by intranasal administration of HDM daily for three days. These CD4 T cells were derived from a T cell receptor (TCR) transgenic mouse specific to Derp-1 (1DER), a class II-restricted immunodominant peptide from HDM. Since 1DER T cells expand in vivo in response to antigen challenge, they comprise a notable fraction of the Th2 population within the lymph node. Spatial mapping in the medLN revealed anatomical organization of T cell zones, B cell follicles, and TB borders, providing the necessary framework for interpreting localizations of 1DER CD4 T cell subsets (**Fig. 6C, Extended Fig. 4A**). Using SpaceTravLR, we sought to identify the strongest spatially dependent ligand-receptor interactions that controlled Th2 identity. Consistent with earlier findings, we recapitulated the importance of IL-2 axis in allergen-specific Th2 cell differentiation while also identifying *Ccr4-*dependent ligand interactions involving *Ccl17*, *Ccl22*, and *Ccl5*^73,74^ (**Fig. 6D**). Interestingly, the distribution of the top interaction scores for these ligands and receptors also exhibited distinct spatial patterns, with Th2 cells closer to the central tissue region showing the highest scores. Specifically, the highest scoring interaction was for *Ccr4*-*Ccl5* in Th2 cells that were localized in the central medLN region, which mirrored the spatial gradient that transitioned from T cell zone to B cell follicles (**Fig. 6E, Extended Fig. 4B**).

**Figure 6:**
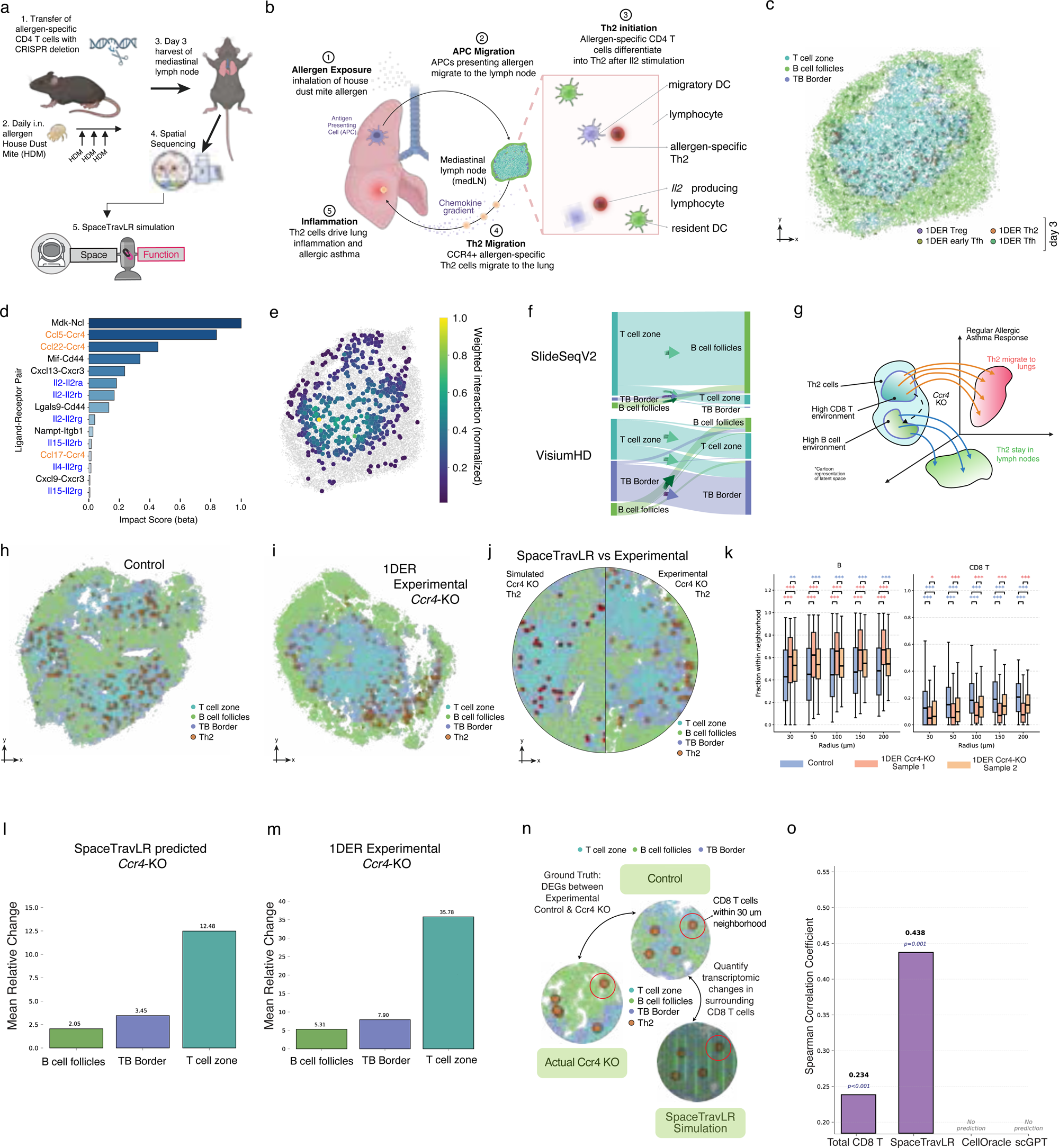
Experimental validation of SpaceTravLR predictions for the spatial relocation of Th2 cells following Ccr4 knockout in murine lymph node. **A.** Schematic of murine model of allergic asthma. CD4 T cells were derived from a T cell receptor (TCR) transgenic mouse specific to Derp-1 (1DER), a class II restricted immunodominant peptide from House Dust Mite (HDM). These cells can be injected directly into the mouse through the tail vein or first altered using CRISPRi to induce *Ccr4* KO. Adoptive transfer is followed by daily intranasal administration of HDM. After 5 days, the medLN is harvested and spatially sequenced with VisiumHD, which can then be used to train SpaceTravLR. **B.** Schematic of Th2 lung-lymph migration patterns. Dendritic cells (DCs) in the lung activated upon encountering HDM migrate to the lymph, where they in turn activate the antigen-specific CD4 T cells through the *Il2*-signaling axis. Activated cells can differentiate into Th2 cells and migrate to the lung and cause inflammation. **C.** Spatial organization of adoptively transferred cells and the anatomical regions (T cell zone, TB-order, B-cell follicles) in the publicly available Slide-seqV2 mouse lymph node harvested 3 days after adoptive transfer. **D.** Ligand-receptor interactions with the highest SpaceTravLR coefficients. (interaction pairs with the C3 gene are not shown in this figure to focus on non-complement based immune interactions). **E.** Th2 cell coefficients for the *Ccl5*-*Ccr4* interaction display the highest importance in the T cell zone which diminishes toward the B-cell follicles. **F.** SpaceTravLR simulated Th2 cell-specific *Ccr4* KO causes Day 3 T-cell zone Th2 cells to become transcriptionally like those in the B-cell follicle. **G.** Novel intra-lymph migration patterns of Th2 cells as uncovered by SpaceTravLR. 1DER Th2 cells typically migrate to the T-cell zone of the lymph node, where the majority of the CD8 T cells reside. *Ccr4*-KO Th2 cells fail to migrate to these regions. **H.** Control and **I.** Experimental *Ccr4*-KO 1DER VisiumHD samples harvested 4 days after adoptive transfer. **J.** SpaceTravLR simulated *Ccr4* KO highlighting the Th2 cell locations (left) vs experimentally observed locations (right). Simulated Th2 cells are shown in red and experimentally observed Th2 cells are shown in orange. **K.** The proportion of B-cells and CD8 T cells within 30, 50, 100, 150, and 200um of the Th2 cells. Across all radii, control Th2 cells are surrounded by significantly more B cells and fewer CD8 T cells compared to both knockout samples. **L.** SpaceTravLR pseudo-bulked transcriptomic change from Th2 cell-specific *Ccr4* KO of the cells within the T cell zone, B cell follicles, and TB borders. **M.** Observed pseudo-bulked transcriptomic change from 1DER cell-specific *Ccr4* KO of the cells within the T cell zone, B cell follicles, and TB borders. **N.** Schematic demonstrating the analysis of CD8 T cell DEGs within a 30μm radius of Th2 cells following *Ccr4* knockout. We identified DEGs by comparing the control sample to both knockouts, selecting genes with consistent changes and absolute log2 fold change >0.5. **O.** Overall transcriptomic difference in each anatomical zone from simulated *Ccr4* KO in control Th2 cells. Overall transcriptomic differences in each anatomical zone between the real VisiumHD control and knockout samples.

To better elucidate the role of *Ccr4* in the context of functional microniches, we applied *in-silico* deletion of *Ccr4* using SpaceTravLR and compared it to CRISPR/Cas9-mediated experimental knockout in 1DER Th2 cells. This provides a very rigorous benchmark of the predictions and novel inferences of our method as the KO is limited to a spatial-niche-restricted cell population. Across both Slide-seqV2 and VisiumHD datasets, our model predicted that *Ccr4* deletion would cause antigen-specific Th2 cells in the T cell zone to become transcriptionally more similar to cells at the TB border and B cell follicles (**Figs. 6F, 6G**). While *Ccr4* is known to promote migration of Th2 cells to the lung, the impact of *Ccr4* deletion on its localization in the lymph node had not been described. SpaceTravLR’s simulated knockout demonstrates high concordance with the observed experimental effect on the Th2 cells across the three zones (**Figs. 6H-J and Extended Figs. 4C-E**). Neighborhood analysis revealed significant differences in the cellular composition surrounding the Th2 cells between the control and knockout cells. In both knockout lymph nodes, Th2 cells were surrounded by a significantly higher proportion of B cells and fewer CD8 T cells, which primarily reside in the T-cell zone. (**Fig. 6K and Extended Figs. 4F, 4G)**. Together, these results demonstrate that our model’s computational predictions can recapitulate in-vivo outcomes. In addition, using SpaceTravLR, we also examined the overall impact of *Ccr4* deletion in the overall local tissue microenvironment. The strongest effects were predicted in the T cell zone, and this was consistent with the actual experimental data (**Figs. 6L, 6M, Extended Fig. 4H**).

We next asked whether SpaceTravLR could capture how antigen-specific (1DER) *Ccr4*-deficient T cells influenced neighboring CD8 T cells (not antigen-specific) in the local microenvironment. Specifically, we compared DEGs in CD8 T cells that were located within 30μm of Th2 cells between control and *Ccr4* knockout samples (**Fig. 6N**). While no existing computational method can fully disentangle these complex relationships, SpaceTravLR produced predictions that closely align with reported experimental observations (**Fig. 6O**). SpaceTravLR’s predictions matched the true local changes within the 30um neighborhood of Th2 cells even more closely than observed changes in the total CD8 T cell pool measured in the experimental sample, indicating that our model successfully captures nuanced, spatially dependent effects of *Ccr4* deletion in the local microenvironment (**Fig. 6O**). These findings suggest *Ccr4* deletion on HDM-specific CD4 T cells may disrupt centrally located Th2 cells and SpaceTravLR predictions can explain how local perturbations can affect regulation and signaling in microniches that impact actual biological functions.

## Discussion

The ability to mechanistically connect biological function to spatial organization remains a major challenge in understanding multicellular systems. Recent developments in computational approaches can generate predictions on cell-intrinsic perturbation effects, but they are not designed to capture how the effects propagate across cellular neighborhoods. Here, we introduce SpaceTravLR, an interpretable machine learning framework that addresses this gap by integrating spatially aware gene regulatory and signaling (ligand-receptor) networks to explain how molecular perturbations shape both cell intrinsic and extrinsic circuits in tissue microenvironments. In doing so, SpaceTravLR moves spatial transcriptomics from a descriptive framework into one that infers causal relationships, enabling discovery of functional microniches, a localized cell cluster connected by causal regulation rather than spatial proximity or similar transcriptional states. Our model predicts how gene expression changes within individual cells alter their signaling, cell fate, and spatial organization. This multi-scale perturbation framework enables unprecedented mechanistic insight into how genetic changes drive tissue-level phenotypes.

Moreover, through applications across diverse tissue contexts, species, and platforms, SpaceTravLR reveals that the impact of genetic perturbation is both spatially constrained and context dependent. Unlike other single-cell perturbation frameworks, our model learns spatially resolved gene-gene relationships and propagates the effects through both transcriptional and signaling networks. This design allows the model to infer *how* a perturbation within one cell type drive transcriptomics and spatial programming in its local tissue neighborhood. As a result, we can interpret these changes through known biological parameters to enable testable hypotheses that link molecular regulators to spatially resolved functional outcomes. Application of our model was able to reproduce known biology from *Foxo1*-mediated germinal center maintenance to *Tbx6* dependent embryo development. It extended to uncovering previously unrecognized spatial reorganization and retention of *Ccr4*-dependent Th2 cells within the mediastinal lymph nodes of a murine model of allergic asthma. In each case, SpaceTravLR was able to broadly infer how intercellular communication shapes tissue architecture and cellular localization.

We anticipate that SpaceTravLR can serve as a foundation for *in-silico* investigation that charts how gene pathways govern tissue organization and spatial microniches. Future work could combine SpaceTravLR with spatial proteomics, temporal modeling, or lineage tracing to enhance mechanistic insights into how genetic perturbations shape complex multicellular structures.

## Methods

### Spatial data preprocessing

All ST datasets were preprocessed using the python framework before training SpaceTravLR. For each, the scanpy^75^ framework was used to perform cell and gene filtering, library size normalization, and log1p transformation^75^. The log transformed data was also smoothed over the cell type specific gene expression manifold using the MAGIC algorithm^76^. For all datasets, we consider the 5000 highly variable genes computed using the scanpy’s highly_variable_genes function. For published Slide-tags and Slide-seqV2 datasets, we used the author provided annotations. Furthermore, for the Slide-tags tonsil and Slide-seqV2 lymph node datasets, we used BANKSY to refine the clustering into subclusters. In tonsil, we subclustered the GC B cells into GC Dark Zones, GC Marginal Zone and GC Light Zone and non-Tfh CD4 T cells into Th0, Th1, Th2 and Tregs using marker genes from the original Slide-tags paper and the Azumith reference-based mapping from the human tonsil atlas. In the lymph node, we clustered the cells into B zone, T zone, and TB border. We filtered beads based on a minimum of 60 total counts and 30 unique genes. We used Tangram^77^ to align a matched annotated scRNA-seq dataset to the Slide-seqV2 data and obtained a probabilistic mapping of cell type onto the spatial coordinates. To eliminate noisy mappings, we applied a cell-type-specific threshold. Finally, we assigned up to three cell identities to each spatial spot, corresponding to the cell types with the highest mapping probabilities above the threshold, to account for potential multiplets or microenvironmental mixing. Spots with high probability (>0.5) of two cell types were split into identical spots with unique identities at the same location while low probability spots were removed.

### Spatially informed regulatory inference from ST

We first construct a base gene modulatory network (GMN) linking target genes (TG) to a curated set of upstream transcription factors, ligands, and receptors. TFs were obtained from CellOracle’s published database of GRNs while L-R and L-TF pairs were obtained from CellChat^78^ and NicheNet^79^ respectively. As an optional preprocessing step, we seek to initially filter out TF-TG and L-R links that the specific ST dataset has weak statistical evidence for. This preliminary filtering provides SpaceTravLR with a higher-quality foundation network for learning tissue-specific gene modulatory networks by eliminating spurious connections. For TF-TG interactions, we apply CellOracle’s Bayesian Ridge framework and perform one-sample t-tests on the learned coefficients to estimate the probability of each being null. For L-R interactions, we apply COMMOT’s Optimal Transport framework to identify cell-cell communication (CCC) while accounting for higher-order interactions between various ligands and receptors. By permuting the location of each cell and recomputing a null CCC baseline, COMMOT can compute p-values for each L-R term and eliminate non-significant interactions.

### Spatially informed signaling inference from ST

Briefly, for each location *j* we define its spatial neighbors *n* as all locations *i* within a circle centered on location *j* with a predefined radius *r*. For the amount of receptor expressed at each location, we directly use the gene expression since receptors are located on cell surface membranes and aren’t diffusible across space. We next calculate the amount of ligand ℒ that a cell at location *j* receives from all *n* neighboring cells weighted by the physical distances between them transformed by a Gaussian kernel Ω, as proposed by previous methods such as CytoSignal^80^. For simplicity, we denote Ω_(*u*,*v*)_ ℒ_*j*_ as 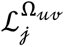. The signaling activity of the k^th^ L-R pair at location *j* can thus be modeled as

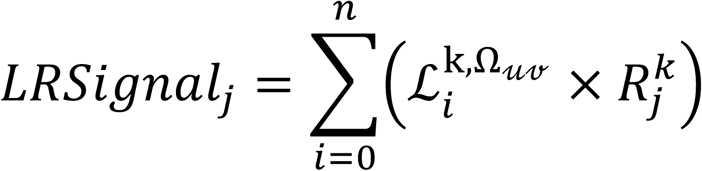

This assumes that the diffusion of ligands from the source cell radially symmetric. In addition to L-R interaction terms, we also incorporate L-TF terms using a similar approach, with the terms collected from the NicheNet database.

### Spatially Weighted Regression

Several methods have been previously proposed for modeling the expression of target genes as a function of its known regulators. Linear models offer an interpretable and easy to train solution for modeling the gene expression profile y in cell type C as a function of m known transcription factors of y.

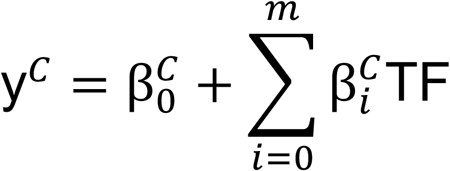

This approach also offers a simple strategy for deriving the direct impact of a particular TF on gene y via the learned coefficients since the derivative 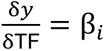. However, to infer functionally distinct microniches across a whole tissue area, we must also model the spatially non-stationary relationships between genes and their upstream modulators - effectively extending β_*i*_ to be a function of location β_*i*(*u*,*v*)_. To this end, SpaceTravLR adopts a Spatially Weighted Convolutional Neural Network Weighted Regression^81^ model to predict the local gene expression of target genes based on the spatially varying expression of regulators, ligands, and receptors.

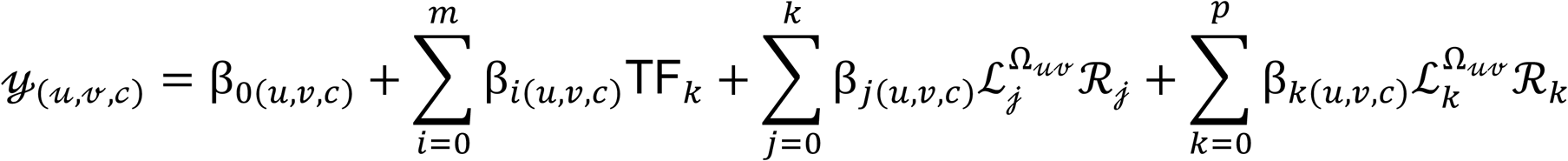

Traditionally, Spatially Weighted Regression (SWR) has been used to estimate the spatially varying relationship^82^ between dependent and independent variables in various research areas such as epidemiology, urban development, air pollution, and agriculture. SWR takes as input a Spatial Proximity Grid (SPG) from which the functional relationship between regression coefficients and spatial distance between samples can be captured.

To construct the SPG, we split the 2D map of cell locations into uniform grids with a predefined resolution (64×64 by default). For each cell i at location (*u*_*i*_, *v*_*i*_), we then compute a spatial distance map 𝒟 between i and the center point of each grid according to

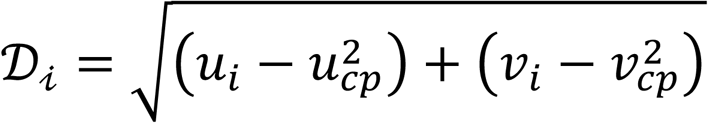

where (*u*_*cp*_, *v*_*cp*_) is the location of center points.

Since user-defined kernel functions typically used in SWR might not be expressive enough to faithfully model complex cell type specific gene expression patterns in biological samples, here we implement a convolutional neural network (CNN) instead. Based on the shared-weight architecture of the convolutional filters that slide along the 2D region, CNNs are able to generate translation-equivariant feature maps. Here we use a 3-layer convolutional network to extract rich spatial information from the input SPG followed by a 2-layer fully connected multilayer perceptron (MLP) with the output dimension of the last layer matching the total number of regression predictors plus the intercept. Optionally, additional spatial information or cell type specific metadata can be concatenated with the flattened spatial features from the CNN to provide more context to the MLP for generating the final output coefficients. Taken together, this SPG → CNN → MLP → β workflow allows us to learn continuous regression coefficients that are functions of location β_*i*_ = *f*(𝒟_*i*_(*u*_*i*_, *v*_*i*_)).

### Filtering ligand-receptor priors with COMMOT

As an optional preprocessing step to filter out uninformative ligand-receptor pairs, we run COMMOT on our input data with default parameters. We separate each pairing with multiple receptor components into individual ligand-receptor component pairs to account for dimerized pairs. We then run COMMOT’s ct.tl.spatial_communication and ct.tl.cluster_communication to compute the p-value by permutating the locations of cell to infer spatial cell-cell communication events. Ligand-receptor interactions that COMMOT identified as not statistically significant are then masked out during training. We choose a looser p-value threshold of 0.3 to identify a broader range of possible ligand-receptor interactions that are further filtered from the spatial information by SpaceTravLR in the next step.

### Fitting a sparse Group Lasso regression model

Next, we build a SWR model in parallel for each gene in the preprocessed ST input dataset. Since genes can be naturally grouped into functional classes (such as transcription factors, ligands or receptors), SpaceTravLR uses a sparse group lasso (SGL) formulation to account for this grouped structure of these predictors. SGL encourages sparsity both between and across groups, eliminating uninformative features to reduce false positive links in the gene-gene interaction network. To improve convergence efficiency and interpretability, we split the coefficients into a non-spatial β^SGL^ and spatial component β^CNN^, learned by the SGL and CNN models respectively such that

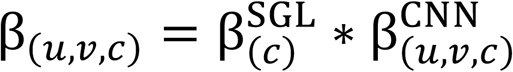

Using A100 Nvidia GPUs, each model takes on average 60 seconds per gene to train for 100 epochs for 10,000 cells. The models are trained to minimize the mean-squared error between the predicted gene expression value and the observed gene expression value at each location in a cell type specific manner.

### Inferring functional microniches driving cell fate

SpaceTravLR can simulate the spatial heterogeneity in perturbation outcomes at a single cell resolution. Our model is designed to simulate the direct impact of gene perturbations on specified cells and the indirect impact on their neighbors. This process consists of 5 key steps: (1) spatial perturbation target selection, (2) intracellular signal propagation, (3) intercellular signal propagation, (4) location specific cell transition estimation, (5) simulated vector field functional alignment.

To illustrate this procedure, here we consider an example gene ℋ modulated by transcription factor 𝒯_1_, ligand-receptor pair ℒ_1_ℛ_1_ and ligand-transcription factor link ℒ_2_𝒯_1_. In spatial transcriptomics data, the gene expressions vary as a function of spatial coordinate (*u*, *v*).

The SWR regression model is thus setup as follows:

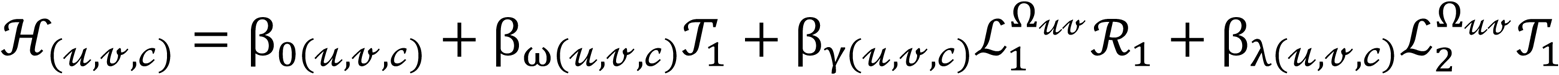

Where 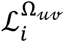 represents the total amount of ligand ℒ_*i*_ received at location u,v

### Spatial perturbation target selection

This step selects specific cells and specific genes to perturb. SpaceTravLR inference requires three inputs:

1. a list of one or more genes to perturb,
2. a list of the desired final expression value for each specified gene
3. a list of one or more cells to target in the given ST data

To prevent out-of-distribution inference, we restrict the desired gene expression values to the minimum and maximum observed values in the training data.

### Intracellular signal propagation

This step computes the transcriptome wide shift in gene expression within each perturbed cell. Modeling the impact of perturbing gene 𝒯_1_ on gene ℋ requires estimating the partial derivative 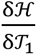. Since SpaceTravLR uses a linear model, this derivative is directly equal to the learned spatial coefficients.

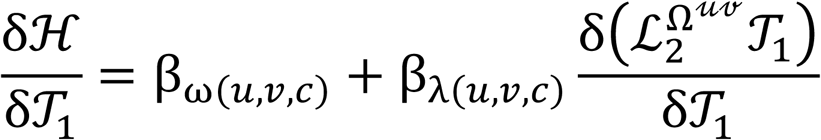

For the ligand terms, we need to compute the partial derivatives by splitting up the ligand interaction coefficient. Assuming that changes in ligand expression and in the paired receptor expression or TF expression are independent, then by the product rule:

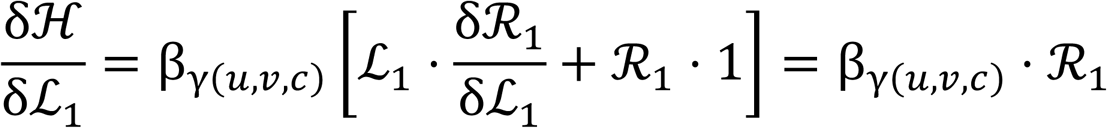

The impact of perturbing ℒ_1_ is thus proportional to the expression of its receptor pair ℛ_1_. Similarly,

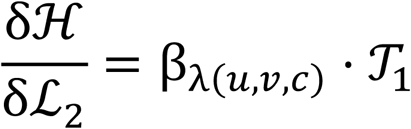

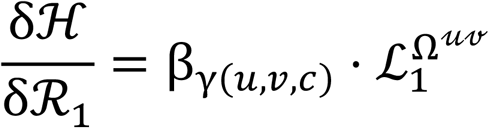

and

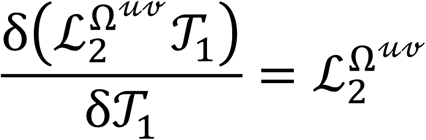

Finally,

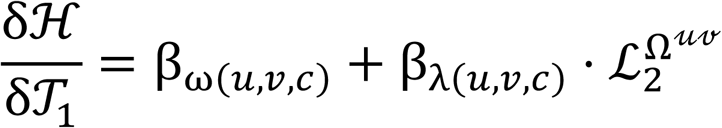

Therefore, the response Δℋ to a perturbation Δ𝒯_1_ at location (*u*, *v*) on cell of type c is given by

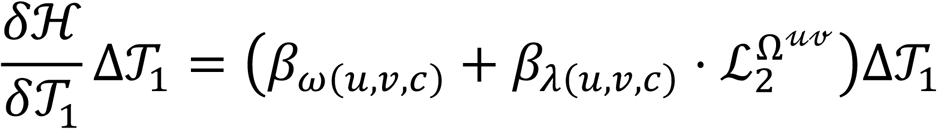

For in-vivo experiments, combinatorial in-vivo perturbations simultaneously target multiple genes, with each gene’s distinct transcriptomic effects propagating through regulatory networks to produce collective changes in cellular phenotype. To accurately reflect the experimental perturbations mechanistically, for combinatorial perturbations, we combine the individual perturbations such that double perturbation of 𝒯_1_and ℒ_1_ is given by

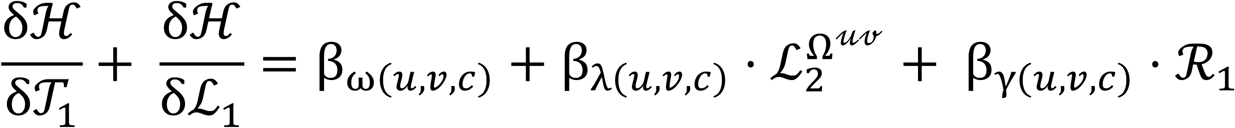

For each possible cell type in a dataset with n cells and g genes, we generate a sparse tensor Θ ∈ ℝ^*n*×*g*×*g*^ where each value represents the gene-gene coefficient as computed above. A second matrix Δ𝒢 ∈ ℝ^*n*×*g*^ is also generated consisting of zero values except for the perturbation gene targets in the specified cells which are set to the shift in expression required to achieve the desired final expression value. To simulate the full impact of a perturbation on all genes, we implement a fast and efficient vectorized operation in PyTorch using sparse tensors.

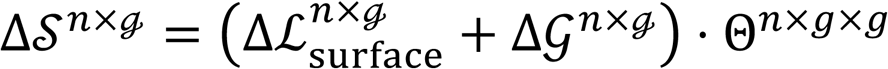

Where Δℒ_surface_ represents the change in ligand signaling received on each cell’s surface at their location and Δ𝒮 represents the transcriptome wide shift in gene expression across all cells.

### Intercellular signal propagation

This step computes the shift in cell-cell signaling and the downstream consequences on gene expression. To compute Δℒ_surface_ we first establish a steady state surface ligand signaling ℒ_baseline_ using the Gaussian kernel Ω.

Then,

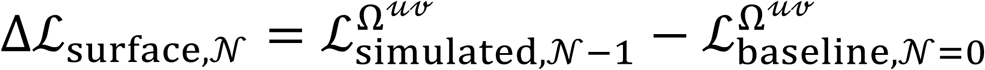

where N is the iteration.

Since a perturbed gene can have second order effects on other genes that it modulates, SpaceTravLR iteratively perturbs genes that are up to 4 nodes away in the network using the chain rule. For example, if gene ℋ modulates some gene ℱ, with 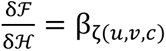, then using the chain rule the response Δℱ to a perturbation Δ𝒯_1_ at location (*u*, *v*) on cell of type c is given by

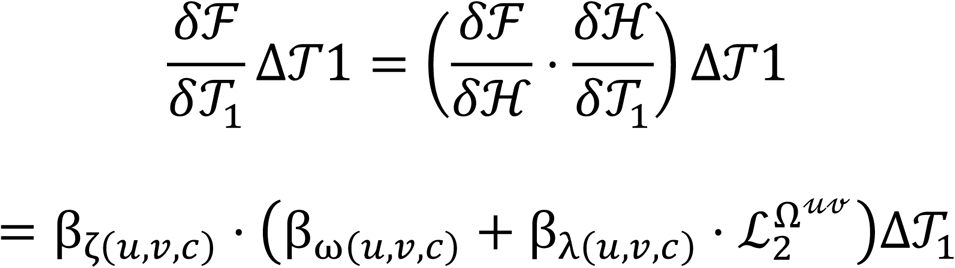

The final simulated output gene expression matrix Δ𝒮 is thus iteratively computed as

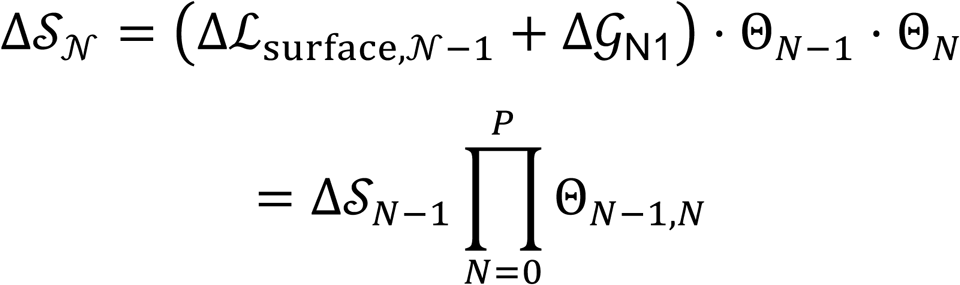

where Θ is recomputed in at the beginning of each iteration (total iterations *N* = 4 by default) using the new perturbed gene expression values given by

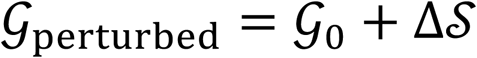

Based on empirical testing, we set the number of propagation steps to 4 (**Supplementary Figs. 8, 9**).

### Location specific cell transition probabilities

This step infers the spatially varying probability of cells transitioning from one state or identity to another. Using the simulated output Δ*S* ∈ ℝ^c×g^, we compute a transition matrix *P* ∈ ℝ^c×c^ where elements *p*_*i*,j_ is the probability that cell i transitions to adopt an identity similar to cell j after a perturbation that shifts its transcriptome from

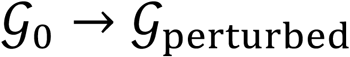

To compute *p*_*i*j_, SpaceTravLR adopts a similar process as CellOracle, using the Pearson’s correlation *d*_*i*_ and *r*_*i*j_

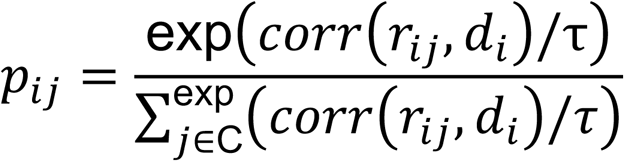

where *d*_*i*_ is the simulated gene expression shift vector Δ*S*_*i*_ ∈ R^1×N^ for cell i and *r_*i*j_* ∈ R^1×N^ is the vector representing the gene expression difference between i and j; *r_*i*j_* = 𝒢_*i*_ − 𝒢_*j*_. The expression is normalized by the Softmax function with default parameter τ = 0.05. For any given cell, the shift in gene expression *d*_*i*_depends on the location of that cell. Unlike previous linear formulations such as CellOracle, SpaceTravLR’s learned β coefficients are functions of cell location (*u*, *v*). As such, we can associate each transition probability *p_*i*j_* with the location of cell i; *p_*i*j_* → *p_*i*j_*_|*u_i_v_i_*_, allowing two transcriptomically identical cells to experience different probabilities of transitioning purely depending on their location in the tissue.

### Simulated vector field functional alignment

To qualitatively assess the functional impact of perturbations, we first project the location specific transition matrix *P* onto a reduced dimensional embedding space (UMAP by default) to create *V*_*i*,projected_ = ∑*p_*i*j_V_*i*j_*, where *V*_*i*,projected_ is the cell-identity transition matrix for cell *i* in reduced dimensions, *V_*i*j_* is the vector computed by subtracting the 2D coordinates in the dimensional reduction embedding between cell *i* and cell *j* and *p_*i*j_* is the cell-cell transition matrix computed above. Next, to visualize the cell-cell transitions, we adopt Velocyto’s vector field framework to compute a summarized vector field *v_g*rid*_* where SpaceTravLR’s simulated transcriptomic shift vector for each cell is grouped by grid point using a smoothing Gaussian kernel.

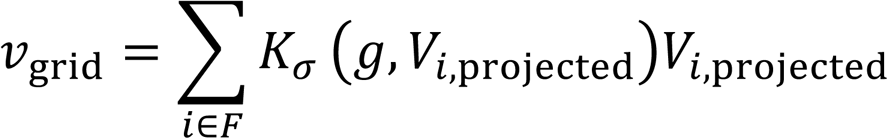

where g denotes the grid point coordinates and F denotes the cells neighboring g. The kernel *K_σ_* is defined by:

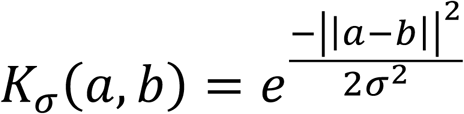

To quantitatively assess SpaceTravLR’s simulations, we then calculated the degree of alignment between our predicted cell state transition and some other independently computed reference score such as differentiation pseudotime, or gene set activity using, the inner product. By calculating the inner product of the simulated perturbation vectors and the gradient vectors of the reference score, we can thus interpret the functional impact of the perturbation - a positive value indicates that both vectors point in similar directions (activating/promoting), a negative value indicates they point in opposite directions (reversing/blocking), and a value of zero indicates orthogonality.

### Benchmarking against scGPT and CellOracle

scGPT and CellOracle are two recent state-of-the-art perturbation prediction methods against which we compare SpaceTravLR. We ran the CellOracle pipeline to perturb transcription factors following their tutorial notebook for Gata1 KO simulation. For scGPT, we downloaded the whole-human pretrained model and finetuned it on our spatial transcriptomics dataset according to their provided tutorial. We extracted gene-level embeddings to construct gene programs according to their GRN tutorial. For each gene, other genes within the same gene program were designated as its modulators, following the linear model format for zero-shot prediction. We then extracted cell embeddings (rather than our spatial-level embeddings) on which we trained our MLP to incorporate cell-level information for fair comparison.

### Transfer learning of spatial coefficients

After learning a spatially weighted linear model for one sample, we can perform predictions on unseen tissue with similar cell type composition. For samples of the same tissue type and similar modality (Kidney XYZeqV2 and VisiumHD), we apply a modified version of SLAT, adapted for aligning heterogenous slices and the learned spatial regression coefficients, enabling us to identify many-to-one and surjective mapping. Multiple cells in the test set may map to the same cell in the train set, and some cells from one sample may not be mapped at all. SLAT incorporates both spatial information and gene expression, accounting for information at multiple scales from individual cells to local niches. To apply a model trained on a higher-resolution modality (Slide-tags), to a low-resolution sample (10x Visium), we align neighborhoods consisting of 5 single-nuclei cells to each 55um Visium spot, which may contain anywhere from 1 to 10 cells. We first run cell2location on the test sample to obtain cell type scores for each spot. For each test spot, we identify the training sample centroid with the most similar neighborhood structure to the cell2location scores based on Pearson correlation. The final coefficients used by the cells in the test sample are the averages of the cell-level coefficients within the matched neighborhood. Once the test samples have been aligned, we are able to use the coefficients to perform perturbations in the same manner as described above.

### Cytope Methodology for XYZeqV2 murine kidney experiments

Kidneys from 8-week-old C57BL/6 female mice were purchased from Envigo Bioproducts and maintained in RPMI 1640 media. The kidney organ was mounted on a microtome (Precisionary Compresstome) and sliced at 300 μm to use on the Cytope microwell array. The microwell array chips were stored at 4°C and equilibrated to room temperature before mounting the kidney tissue slice on top of the array. A digital image was taken to record the tissue orientation after the tissue was pressed onto the Cytope array, secured between the chip and a glass microscope slide. The chip was placed inside a humidity chamber at room temperature and incubated for 30 minutes. After the incubation period, the chip was removed from the humidity chamber and immediately washed in an initial reagent reservoir containing 40 ml of RPMI with 10% FBS which was followed by a second wash in a separate reagent reservoir containing 25 ml of PBS to rapidly dilute and remove any unbound spatial label. The tissue slices were transferred into their respective sample tubes containing 0.5 ml of 2 kU/ml Collagenase IV with protease inhibitor (Millipore Sigma C6079) and dissociated at 37°C for 30 minutes with trituration every 10 minutes. The cells were filtered through a 70 μm cell strainer and washed once with RPMI and 10% FBS. To ensure fluorescence activated cell sorting (FACS) of only the cells that are labeled with the printed antibodies from the array, the cells were stained with 1 μl of PE goat anti-rat polyclonal secondary antibody (Biolegend 405406) in 100 μl of PBS for 30 minutes on ice. The cells were washed twice, with 2 ml of PBS and 10% FBS, and resuspended in a final volume of 0.5 ml of 25 nM Helix NP NIR (Biolegend 425301). All samples were strained through a 35 μm filter before FACS.

### FACS Sorting Methodology for XYZeqV2 murine kidney experiments

To acquire high quality cells for downstream processing, cells characterized as smaller than 4μm and doublets were gated out. Cells that were negatively stained for the viability stain, Helix NP NIR, were gated into a FSC v PE channel plot where cells that were positively labeled for the goat anti-rat polyclonal secondary were sorted into a 1.5 ml Eppendorf tube coated with heat-inactivated FBS.

### 10x Genomics Chromium Platform/Library Preparation for XYZeqV2 murine kidney experiments

Sorted cells were centrifuged and resuspended in phosphate-buffered saline at 1000 cells/ul and loaded on the 10x Genomics Chromium platform and processed using the Next GEM Single Cell 3’ with Feature Barcoding protocol for library preparation. Sample libraries were sequenced on the Novaseq X.

### Adoptive Transfers, CCR4 deletion and HDM immunization in VisiumHD murine lymph node experiments

C57BL/6J (0000664) were purchased from Jackson Laboratories. 1DER mice were previously described^83^. All animals were housed in specific pathogen-free conditions in the Assembly building. All experiments were approved by the University of Pittsburgh Institutional Animal Care and Use Committee. All mice were over 6 weeks of age and both male and female mice were used. Naive CD4 T cells were enriched from 1DER TCR Tg mice using Naive CD4 T cell isolation kit (STEMCELL,19765). For Ccr4 KO by CRISPR– Cas9, naive CD4 T cells were electroporated with murine target or scramble control Single-guide RNA (sgRNA). The detailed protocol using the CRISPR–Cas9 system was previously described^84^. sgRNA targeting murine Ccr4 and scramble control sgRNAs were obtained from Synthego (CRISPR Gene Knockout Kit v2). The Cas9 nuclease V3 was obtained from Integrated DNA Technologies (1081059). The sgRNA–Cas9 RNA complex was introduced into naive CD4 T cells using Lonza 4D-Nucleofector Core Unit (Lonza, AAF-1002B). All deletions were validated by flow. After CRISPR/Cas9 deletion, a total of 1×10^6^ CD4 T cells were injected into congenic recipients. After 24 h, mice were immunized with HDM (25 μg of lipopolysaccharide-low HDM (Stallergenes Greer) in PBS) intranasal to mice anesthetized with isoflurane daily for 3 days. Mice were sacrificed on day 3, mediastinal lymph nodes harvested and fixed in formalin for paraffin embedding.

### VisiumHD protocol and analysis

FFPE sections (10um) were prepared on glass slides, H&E stained and subjected to the VisiumHD protocol per the manufacturer’s recommendations (10x Genomics). Images were acquired with a Leica ImageScope. Libraries were sequenced on an Illumina Nextseq2000 at a depth of 275 million read pairs per capture area. Sequencing files were initially processed by spaceranger (10x Genomics) to align reads to GRCm38/mm10 and mapped with the image. The high-resolution image was aligned using the Visium HD Manual Alignment tool on Loupe Browser. Five matched landmarks per sample within the capture area were selected for the alignment and then algorithmically refined by the software. We then used the Bin2cell^85^ algorithm to reconstruct single cells more accurately by segmenting the H&E images. Finally, we assigned cell type labels to each segmented cell using cell2location.

### VisiumHD Preprocessing Pipeline

All of the newly generated mouse lymph node VisiumHD samples were first processed through SpaceRanger. Using the H&E staining, we applied Bin2Cell to segment the nuclei in the samples. After identifying the locations of the nuclei, the 2um bins were combined to generate the full cell-level transcriptomic profiles, more accurately representing each cell. Since we had 4 lymph node samples per slide, we used Napari to manually select the areas containing each slice. In order to annotate the cells, we applied various methods, including Tangram and RCTD^86^. However, cell2location worked the most accurately based on expert understanding of cell type organization within the lymph node. We utilized the cell2location scores, using the same reference dataset generated from the Slide-seqV2 sample. Due to the sheer size of each lymph node, with around 80k cells each, we randomly divided the cells into 4 groups, running cell2location on each group separately. This approach retains the relative proportions of the cell types within the slice, and is also hypothetically similar to having 4 thinner, adjacent slices. Although we began with 4 slices for each condition, only one control sample and two 1DER *Ccr4*-KO samples had proportions and architectures that matched our biological knowledge. The other slices were dominated by extremely large numbers of B-cells, likely due to the fact that the slices were taken closer to the edge of the lymph node rather than the center. We then continued our pre-processing using only the lymph nodes with representative cell type proportions. Similar to the processing for the other datasets, we used COMMOT to filter the initial number of ligand-receptor pairs. However, due to the computational requirements of optimal transport, we only applied COMMOT to a random subset of 20k cells. For training SpaceTravLR, we used the same 4 Cell2location splits. For network propagation, we restricted TF-gene links to those also identified by CellOracle to prevent spurious associations.

## Data availability

We downloaded the annotated human tonsil Slide-tags dataset from Broad Institute Single Cell Portal under the accession number SCP2169. We downloaded the human melanoma annotated dataset from Broad Institute Single Cell Portal under the accession number SCP2176. The XYZeqV2 murine kidney dataset was generated and annotated in collaboration with SurveyGenomics. We downloaded the annotated Slide-seqV2 murine embryo dataset from the Chan Zuckerberg CELLxGENE portal. Our generated spatial transcriptomics datasets (two mouse kidneys at true single-cell resolution generated by Survey using novel technology, one VisiumHD mouse lymph node in the allergic HDM model, and two VisiumHD mouse lymph nodes with knockout adoptively transferred cells) are available at https://github.com/jishnu-lab/SpaceTravLR

## Code availability

SpaceTravLR code, data, documentation and examples are available on GitHub at https://github.com/jishnu-lab/SpaceTravLR

## Acknowledgements

J.D. was supported in part by NIAID DP2AI164325, NIAID R01AI170108, NIAID U01AI179514 and NHGRI U01HG012041. Y.L. has received funds from the Parker Institute for Cancer Immunotherapy (PICI). A.C.P. was supported in part by NIAID R01 AI153104 and NIAID R01 AI156093. The authors acknowledge the resources and services provided by the Rangos Histology Core Facility, a shared facility in the Rangos Research Center at the University of Pittsburgh. This project utilized the services of the University of Pittsburgh Health Sciences Sequencing Core at UPMC Children’s Hospital of Pittsburgh to perform VisiumHD. This research was supported in part by the University of Pittsburgh Center for Research Computing and Data, RRID:SCR_022735, through the resources provided. Specifically, this work used the H2P cluster, which is supported by NSF award number OAC-2117681 and the HTC cluster, which is supported by NIH award number S10OD028483. Figures 1, 2a, 4a, 5b, 6a,b were created with BioRender (https://www.biorender.com).

**Extended Figure 1: SpaceTravLR recapitulates expected outcomes across diverse perturbation settings and spatial contexts.**

SpaceTravLR’s predictions of the relative impact of lineage defining transcription factors (left), ligands (center), and receptors (right) on the cell types in **A.** Comparison between SpaceTravLR and CellOracle TF KO

SpaceTravLR’s predictions of the relative impact of lineage defining transcription factors (left), ligands (center), and receptors (right) on the cell types in

**B.** murine kidney profiled using XYZeqV2

**C.** human melanoma sample profiled using Slide-tags

**D.** murine lymph node profiled using Slide-seqV2.

**Extended Figure 2: Additional Slide-tags tonsil experiments**

**A.** Schematic of Tfh-DAS

**B.** UMAP projection (left) and spatial layout (right) of cells with GC dark zone B cells colored by likelihood to transition to GC light zone B cells. The degree of impact of perturbation, as quantified by the transition probability, varies spatially. Cells that are transcriptionally more similar can have different transition probabilities because they are in different niches.

**C.** Spatial visualization of GC B cell structure in the WT (left) and CellOracle Foxo1 simulated knockout in GC B cells (right).

**D.** Quantification of the CellOracle predicted transitions from *Foxo1* knockout in GC B cells. *Foxo1* knockout causes a notable portion of GC light zone cells to become GC marginal zone cells, which does not align with known biology. Distribution of the simulated cell-level log2 fold changes predicted by

**E.** SpaceTravLR and **F.** CellOracle. Only the genes with the greatest log fold changes predicted by CellOracle are shown. Only the top 13 genes show non-negligible changes.

**G.** Distribution of predicted log2-fold changes per cell for genes downstream of Foxo1 without cell-cell communication (CCC) terms, achieved by removing ligand-receptor and ligand-TF associations from SpaceTravLR’s linear model.

**H.** Gradient tracking through the gene modulatory network to investigate *Foxo1*’s impact on *Cxcr4*, displaying the top 5 most influential genes at each network layer. The presence of receptors *Cxcr4* and *Cd74* and ligand *Mif* in this pathway demonstrates that CCC is necessary for accurate prediction of *Foxo1*’s effect on Aicda.

**I.** Treg-like cells in the tonsil demonstrate overall downregulation of Treg-associated genes including *Foxp3* and *Ccr4*. Instead, markers associated with Th1 cells, including *Cxcr3* and Ifng are upregulated. Genes not associated with the alternative Treg/Th1 pathway from *Foxo1* knockout show little to no change.

**Extended Figure 3: Examination of alternative paracrine signaling pathways and microniches in XYZeqV2 murine mouse kidney.**

**A.** Top 5 most differentially expressed genes distinguishing hypertonic and regular macrophages identified in the XYZeqV2 dataset.

**B.** UMAP projection (left) and spatial layout (right) of cells with hypertonic macrophages colored by likelihood to transition to regular macrophages. The degree of impact of perturbation, as quantified by the transition probability, varies spatially.

**C.** UMAP projection (left) and spatial tissue layout (right) of all cells (top) and just myeloid cells (bottom) colored by microniche assignments identified by BANKSY clustering.

**D.** UMAP projection (left) and spatial tissue layout (right) of myeloid cells colored by microniche assignments identified by COVET.

**E.** Gene expression histogram of the ligand component *Tgfb2* and its receptors Tgfbr1 and *Tgfb3*. While *Tgfb2* is primarily expressed by the three cell types: endothelial, mesenchymal stromal, and epithelial cells, the receptors are most highly expressed by myeloid cells.

**F.** The specific genes affected and the extent to which they are affected in myeloid cells are dependent on which of the three cell types was perturbed.

**Extended Figure 4. Additional analyses of perturbation experiments and SpaceTravLR results.**

**A.** Cell type composition of the cell types annotated in the publicly available Slide-seqV2 murine lymph node harvested 3 days after adoptive transfer.

**B.** Spatial distribution of the learned Th2 cell coefficients for the interactions *Ccl5*-*Ccr4* and *Ccl22*-*Ccr4* interaction in Slide-seqV2 murine lymph node.

**C.** Spatial organization of the cell types in the novel VisiumHD murine lymph nodes harvested 4 days after adoptive transfer.

**D.** Experimental pipeline of control and 1DER *Ccr4* knockout experiment. Mean Fluorescence Intensity (MFI) demonstrates successful knockout.

**E.** Spatial organization of the Th2 cells in the second 1DER *Ccr4* knockout sample. Th2 cells are concentrated at the edges of the T cell zone.

**F.** Cell type composition of the 3 novel VisiumHD murine lymph nodes.

**G.** Co-occurrence score of the Th2 cells with other cell types at different radius distances. The Th2 cells exhibited differential co-occurrence scores between control and knockout samples, further corroborating the observation of altered Th2 organization within the lymph node. Plots were generated using Squidpy’s co-occurrence probability computation.

**H.** Observed pseudo-bulked transcriptomic change from 1DER cell-specific *Ccr4* KO of the cells within the T cell zone, B cell follicles, and TB borders in second *Ccr4*-KO experimental sample.

## Supplementary information

The file contains Supplementary Figures 1-12

