## Extended Figures for "Characterizing spatial functional microniches with SpaceTravLR"

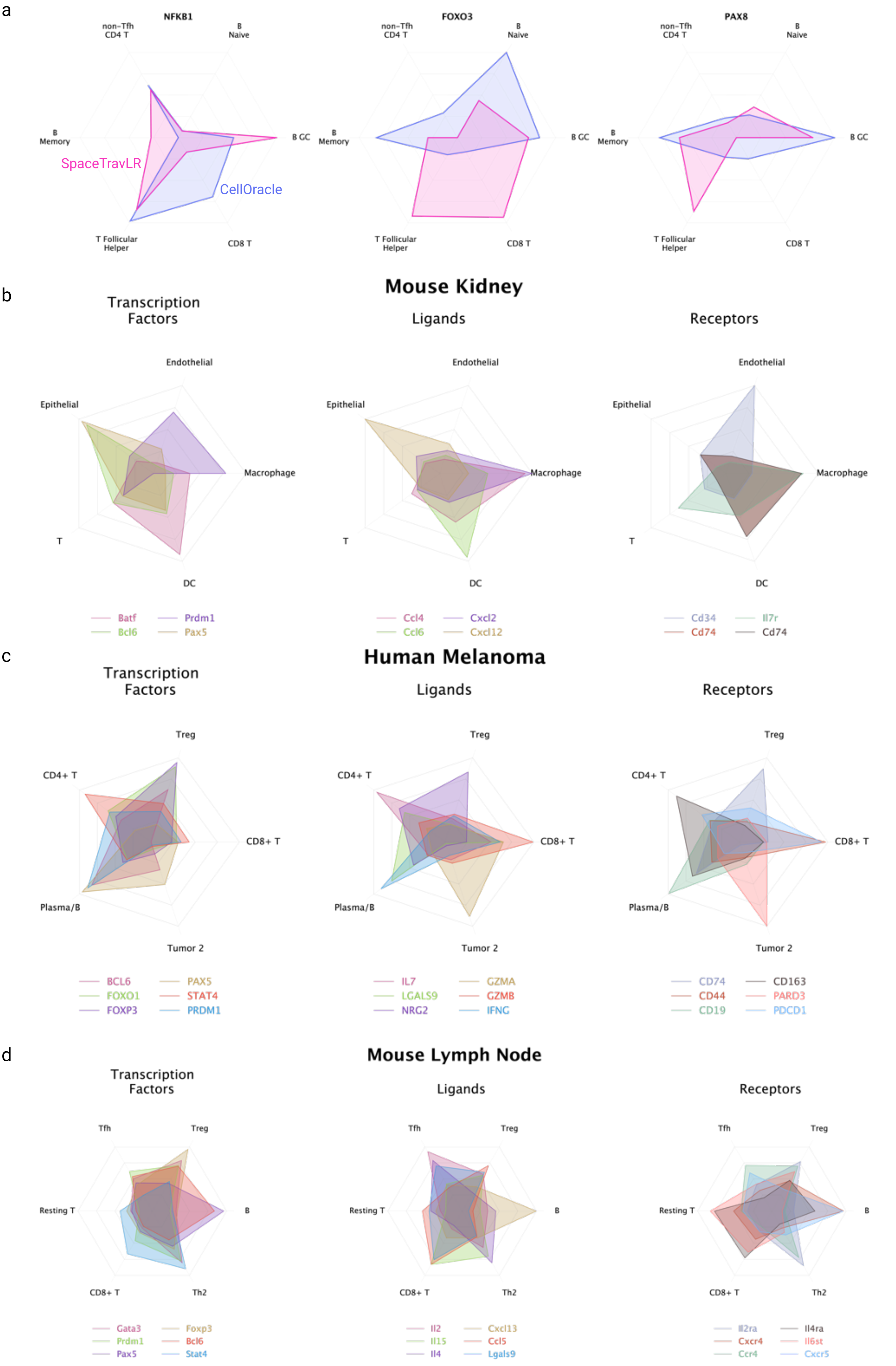


Extended Fig. 1

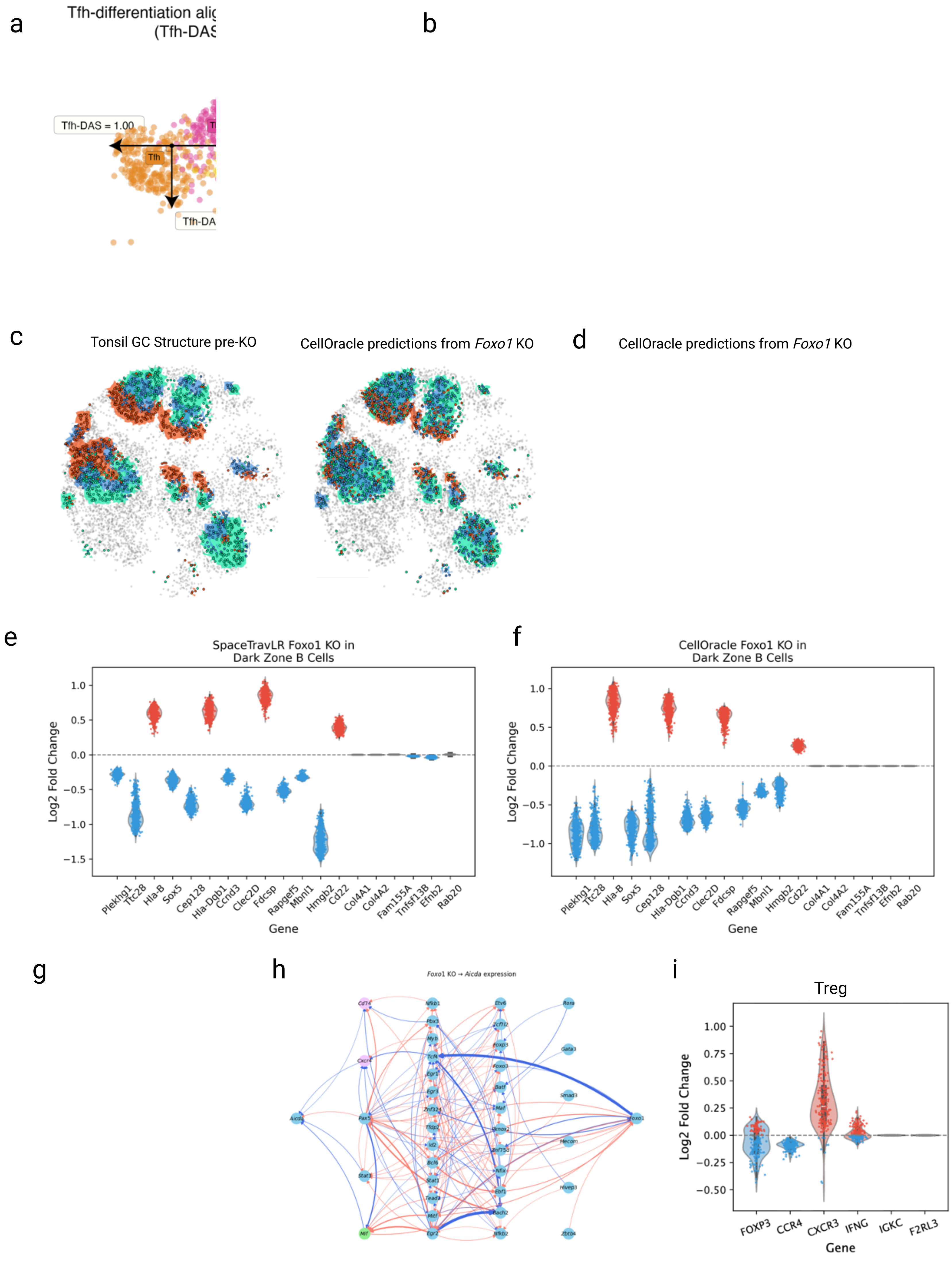


Extended Fig. 2

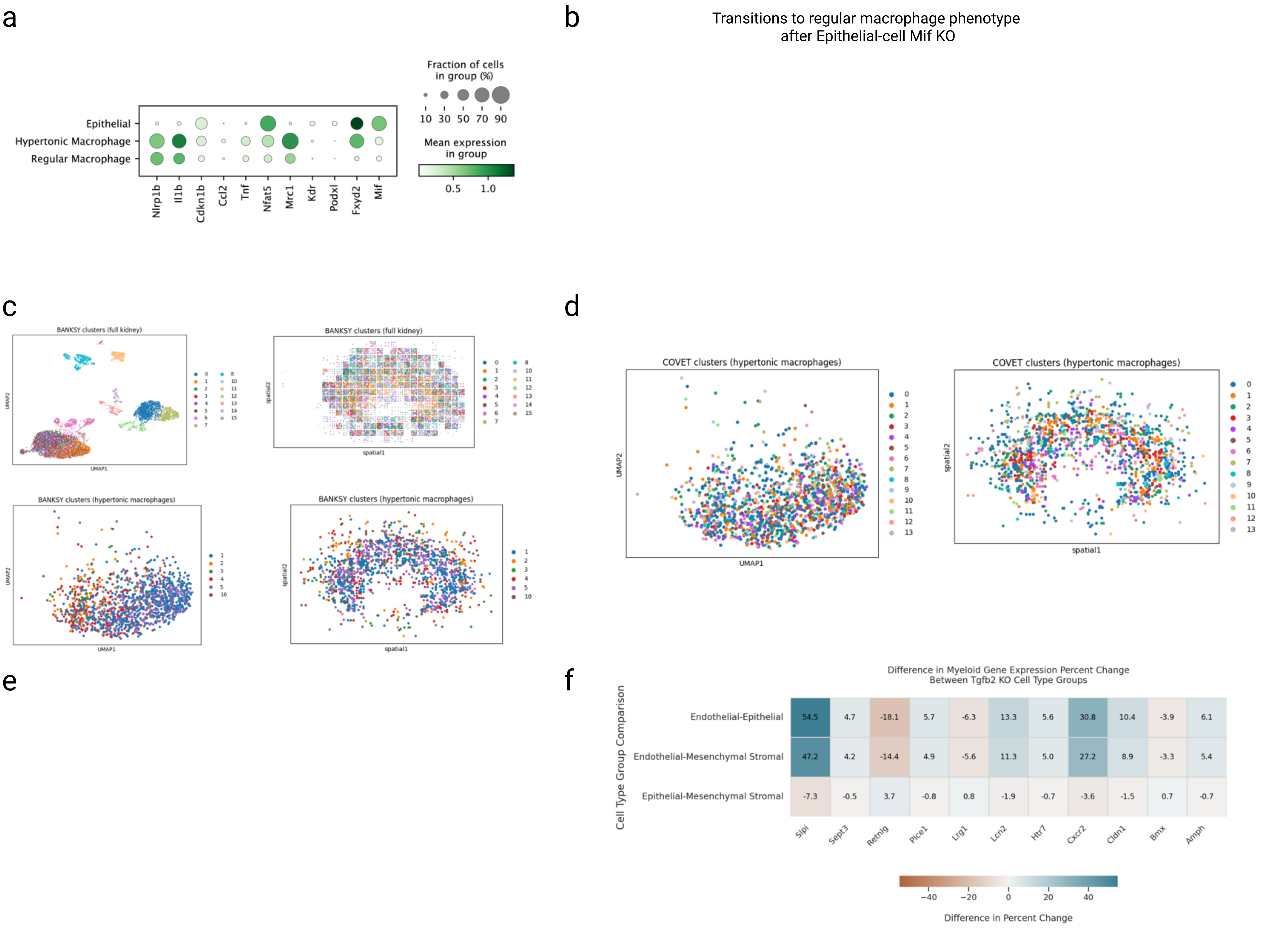


Extended Fig. 3

a

Slide-seqV2

2.8%
8.0%
9.4%
53.2%
16.5%
9.1%
1.0%

b

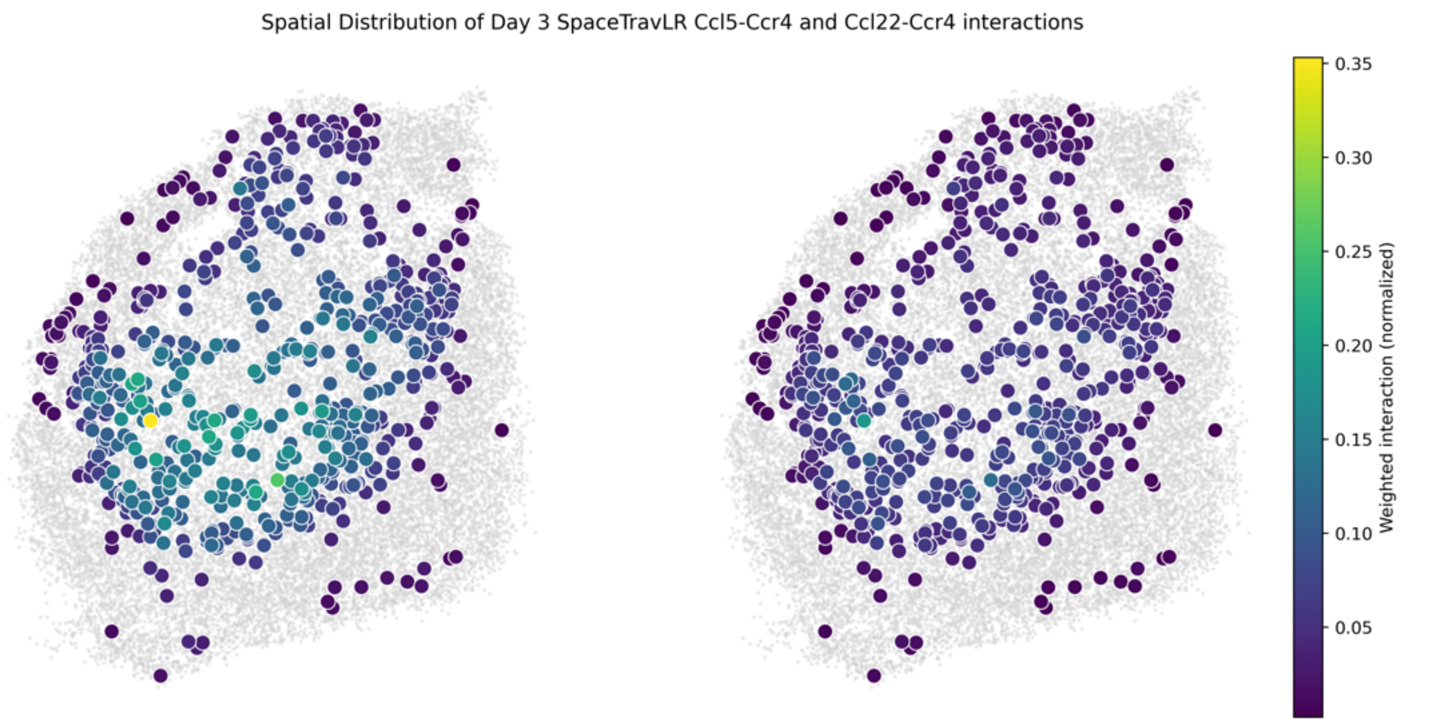


c

Control

1DER *Ccr4*-KO
Sample 1

1DER *Ccr4*-KO
Sample 2

d

e

f

Control

0.4% 1.8% 0.5% 1.2%
0.3%
2.0%
7.8%
10.2%
75.7%

1DER *Ccr4*-KO
Sample 1

0.4% 1.7% 0.4% 0.8%
0.4%
1.9%
10.5%
13.9%
70.0%

1DER *Ccr4*-KO
Sample 2

0.4% 1.7% 0.5% 1.7%
0.3%
13.8%
2.6%
14.0%
65.0%

g

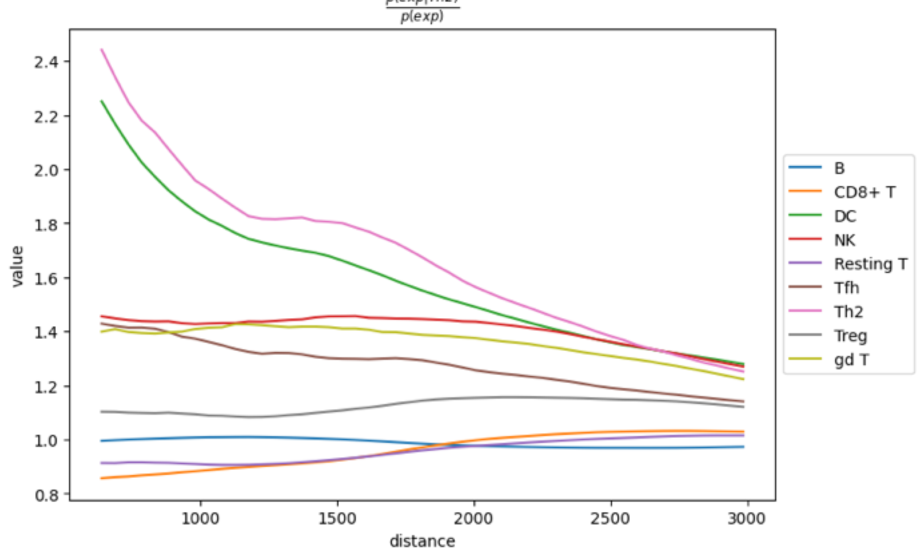


h

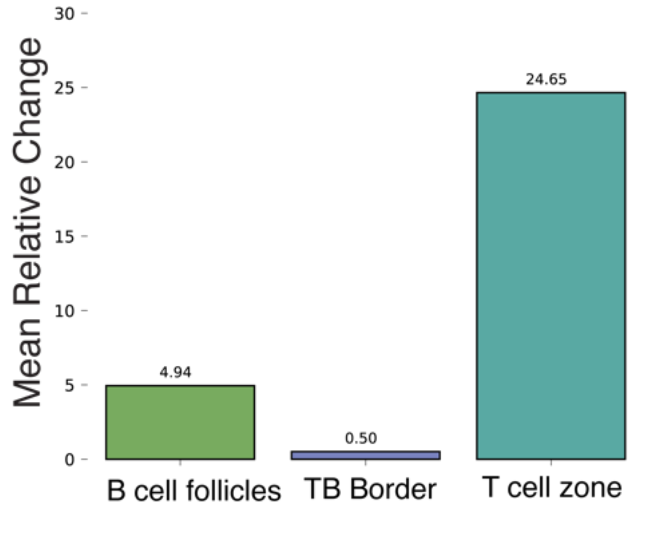


Extended Fig. 4
