## Supplemental Figures for "Characterizing spatial functional microniches with SpaceTravLR"

### Supplementary Figure 1. Impact of radius parameter on LR modeling

a

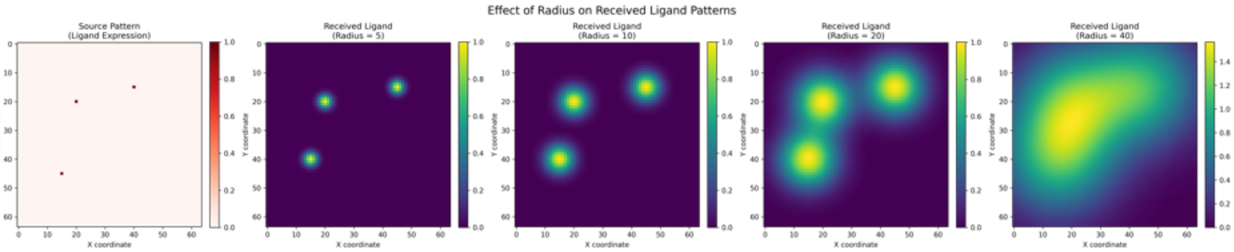


b

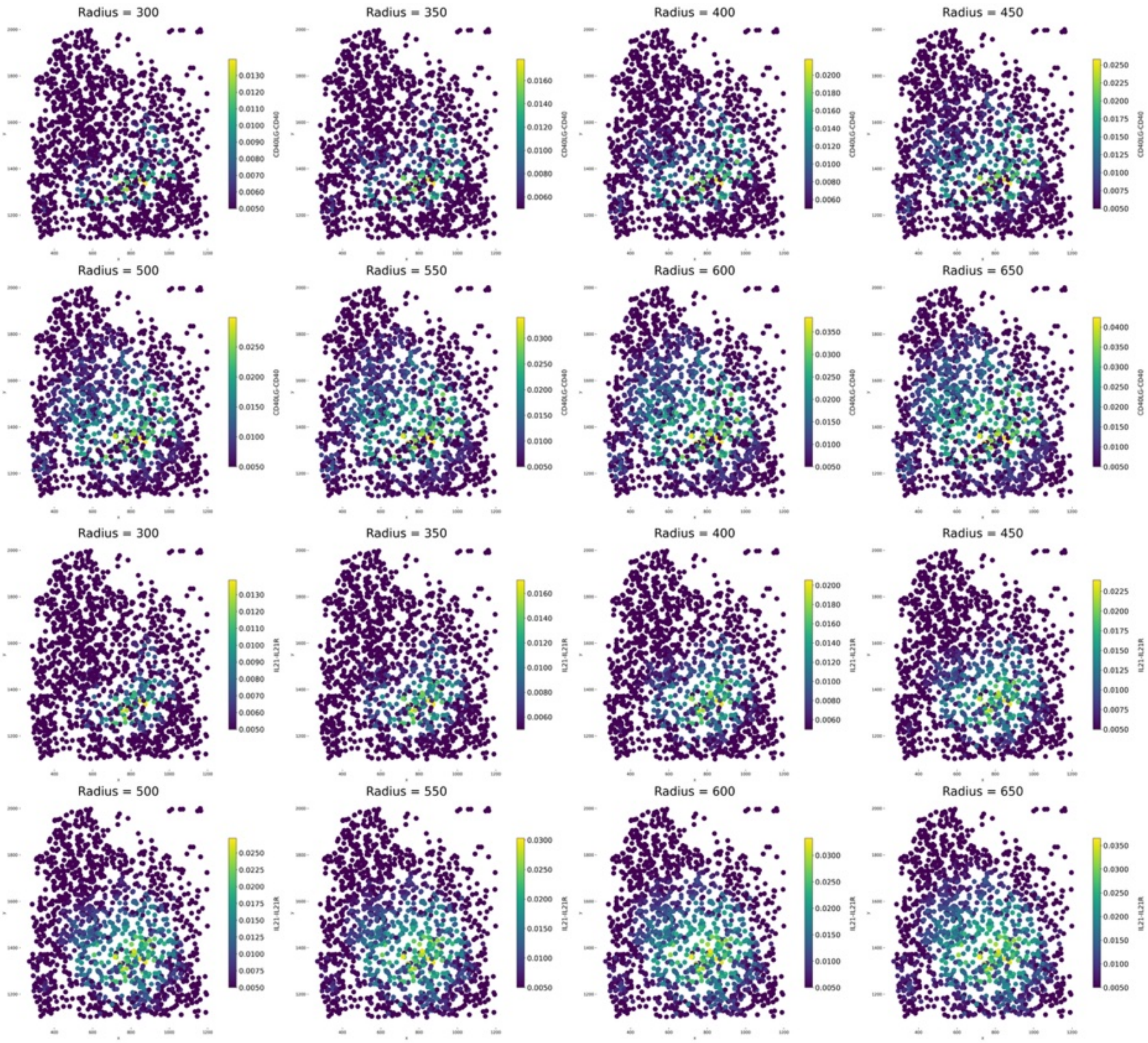

C

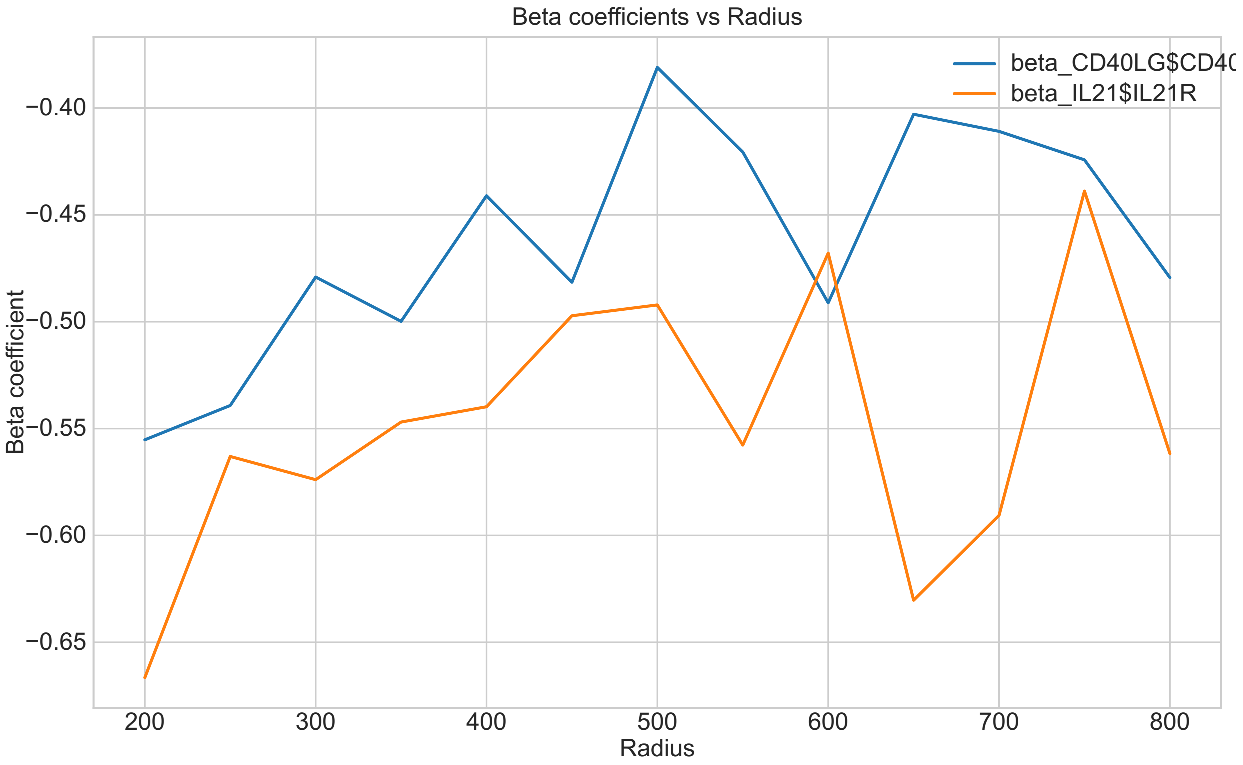


A. Simulated ligand diffusion under gaussian kernel. Increasing the radius increases the total amount of ligand received around the cell symmetrically. B. Effect of radius parameter on effective LR score. The LR score represents the total amount of ligand receive SpaceTravLR defaults to 400-600 microns to avoid missing important interactions, with higher radius favoring poorer quality datasets where ligand expression might not be well captured. C. Increasing the radius parameter impacts the learned regression coefficients, but values converge at 400+ microns. Much larger values lead to unstable training

Supplementary Figure 2. SpaceTravLR intrinsic and extrinsic network propagation

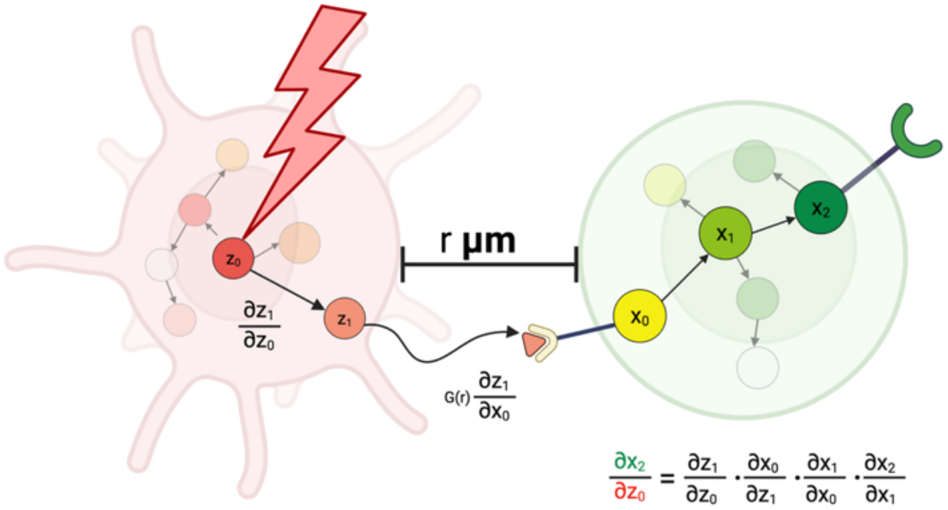


A. Schematic of partial derivatives and chain rule used to propagate signal within and between cells. Here, transcription factor $Z_0$ in a regulator of ligand $Z_1$. $Z_1$ interacts with a receptor $X_0$ which modulates the downstream transcription factor $X_1$, and $X_1$ in turn regulates another receptor $X_2$.

#### Supplementary Figure 3. SpaceTravLR accurately recovers spatially dependent coefficients on simulated data

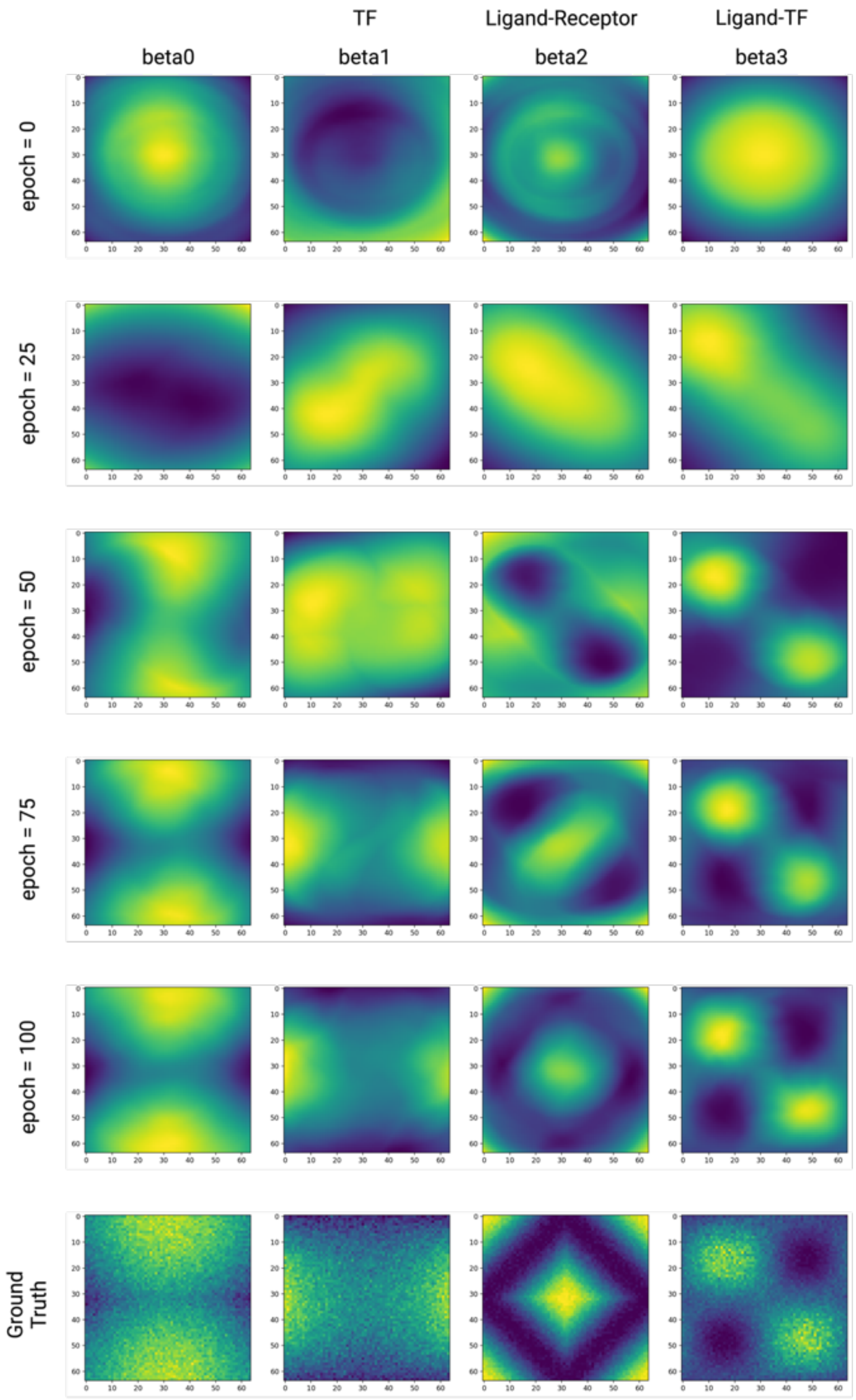


SpaceTravLR's estimation of spatially distinct coefficients of simulated data. beta0 corresponds to baseline effects, beta1 to transcription factor terms, and beta2 and beta3 for the ligand-receptor and ligand-transcription factor interaction terms. SpaceTravLR's vision model successfully recovers all spatially distinct coefficient patterns after 100 epochs of training. (last row) Ground truth synthetic spatially varying coefficients.

Supplementary Figure 4. Spatial coefficients learned by SpaceTravLR in GC B cells

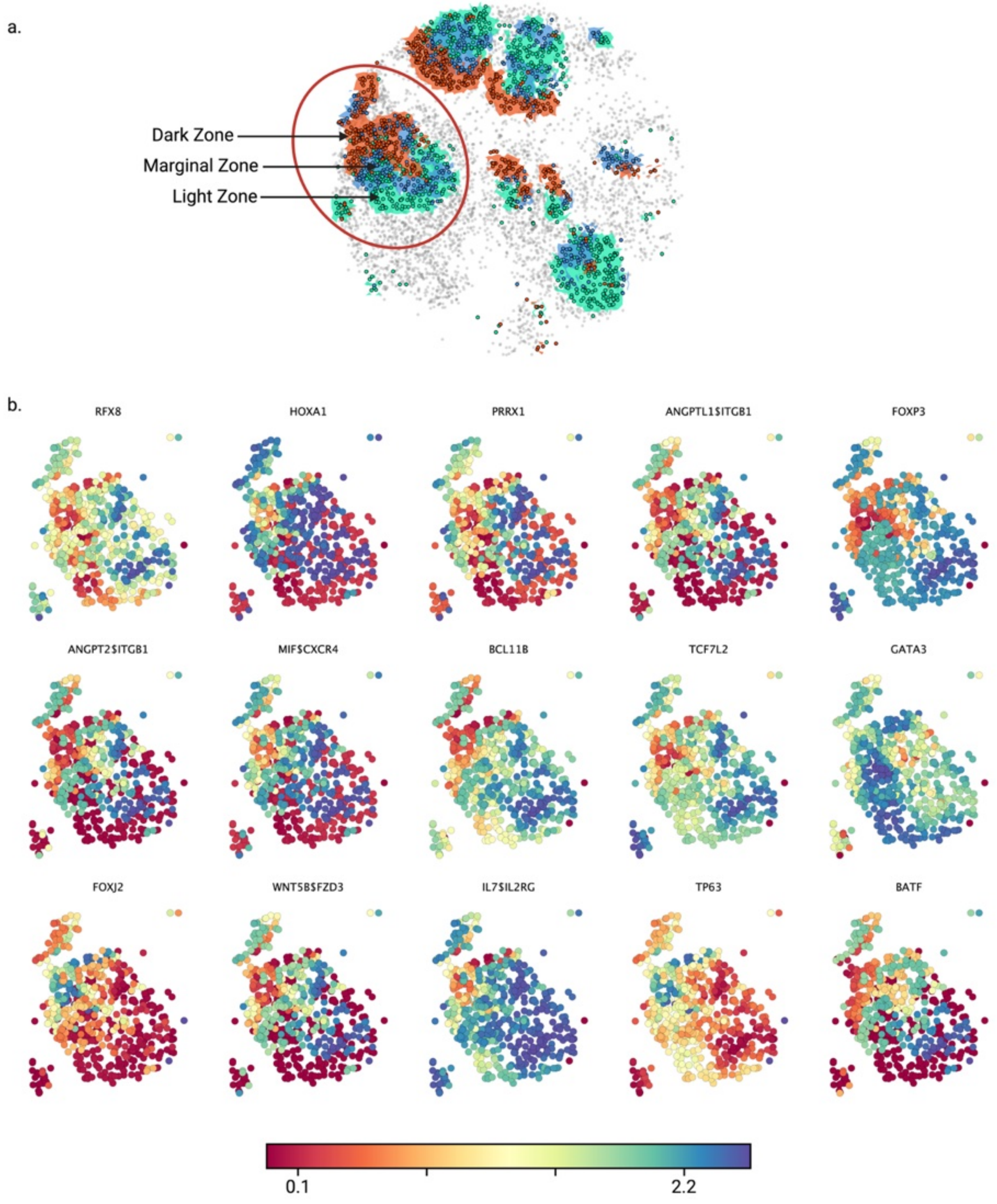


A. Zoomed in section of GC B cells in Slide-Tags tonsil selected for spatial visualization.
B. The modulator terms for the selected GC B cells FOXO1 are spatially distinct, many of them mirroring the spatially distinct zones within the germinal center.

#### Supplementary Figure 5: Comparison of SpaceTravLR model using ViT vs CNN

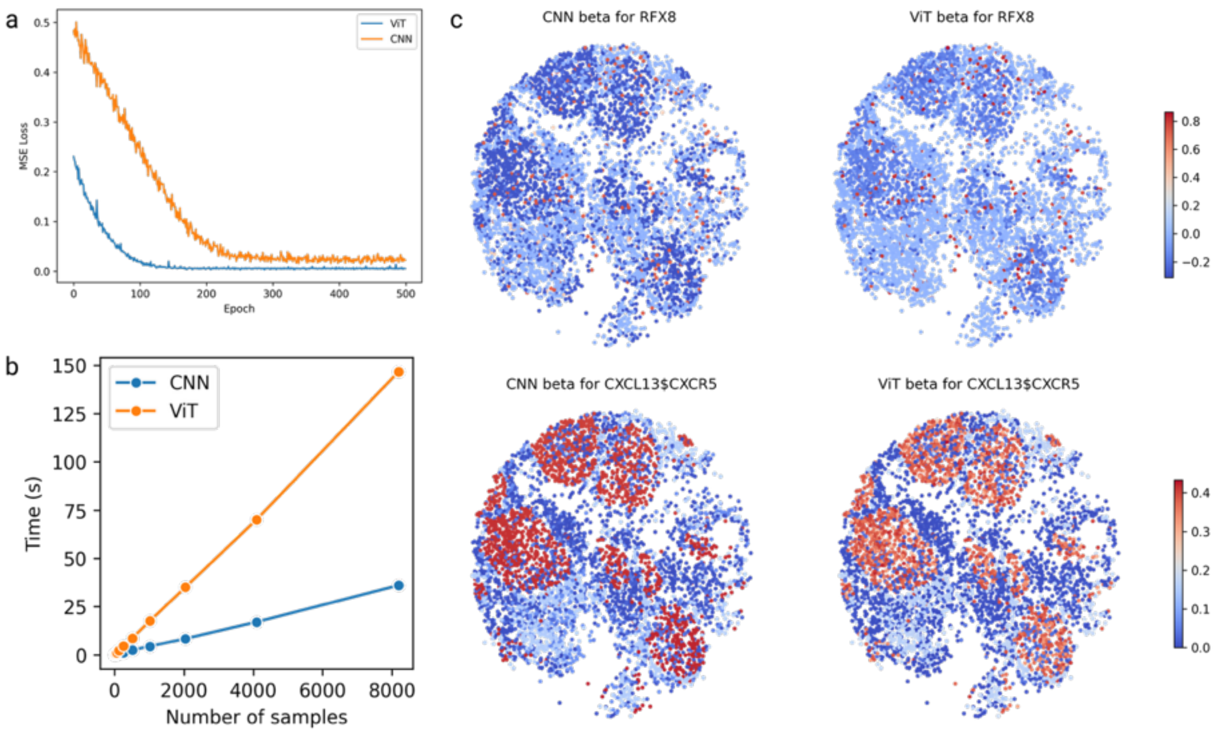


A. Comparison between Convolutional Neural Network (CNN) and Vision Transformer (ViT) training performance, with both achieving similar mean squared error (MSE). B. Scalability of CNN and ViT with increasing sample size. C. Comparison of learned spatial coefficients between CNN and ViT, showing very similar values at convergence.

Supplementary Figure 6. Comparing different regression backbones in fitting the linear model of SpaceTravLR

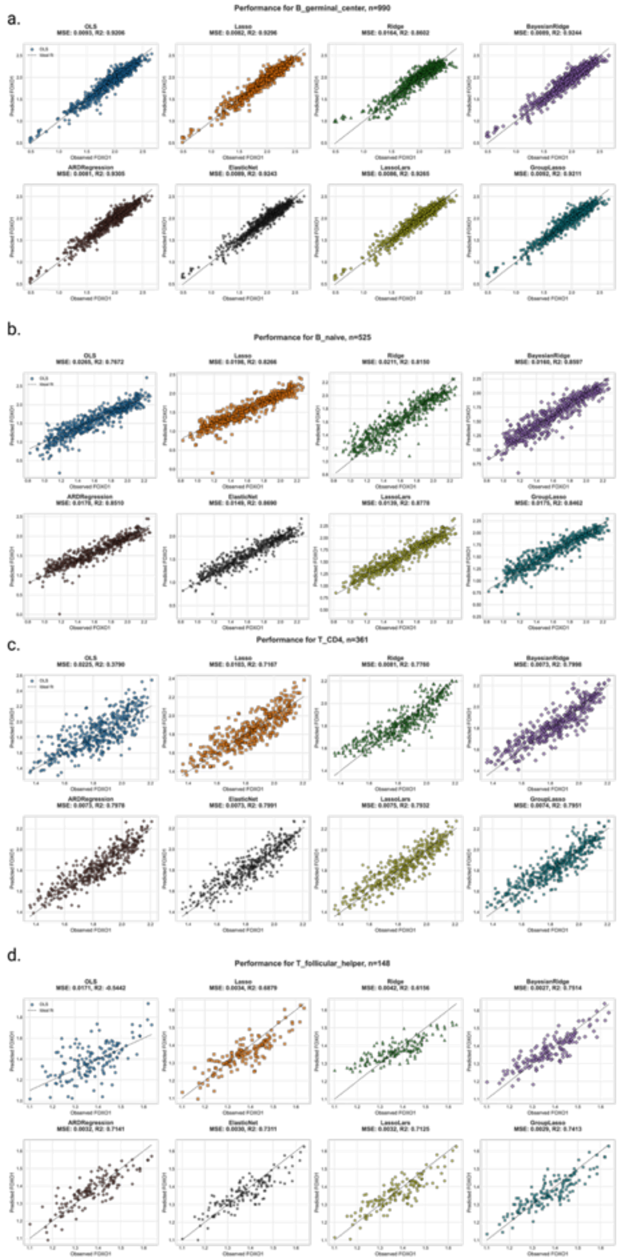


Comparison of 8 different backbones' ability to estimate true FOXO1 expression in A. Germinal center B cells, B. naïve B cells, C. non-Tfh CD4 T cells, and D. Tfh cells in the Slide-Tags human tonsil sample. The performance metrics mean squared error (MSE) and $R^2$ reveal similar performance across model architectures for larger cell populations, such as the B cells. However, Bayesian Ridge and Group Lasso perform the best for smaller cell populations such as in the Tfh cells.

### Supplementary Figure 7. Comparison of model coefficients estimated from different regression backbones

a

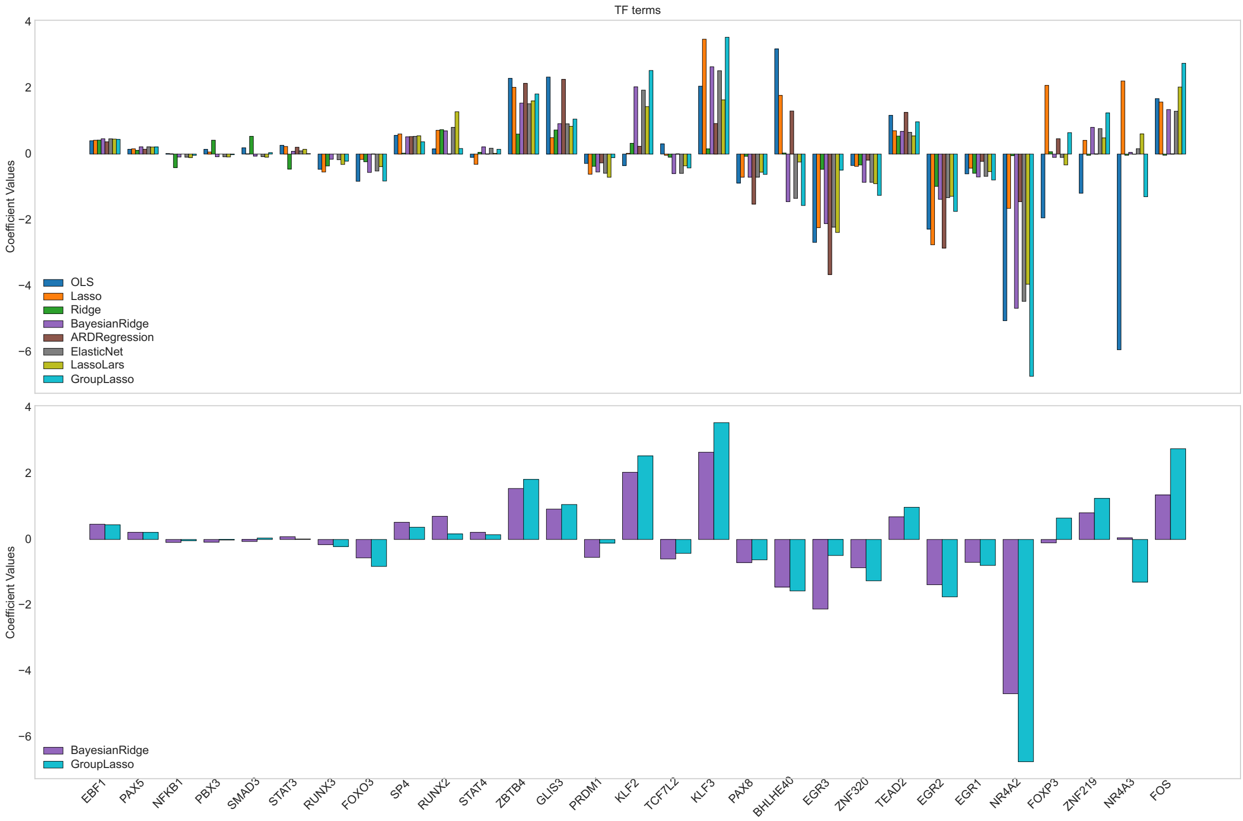

b

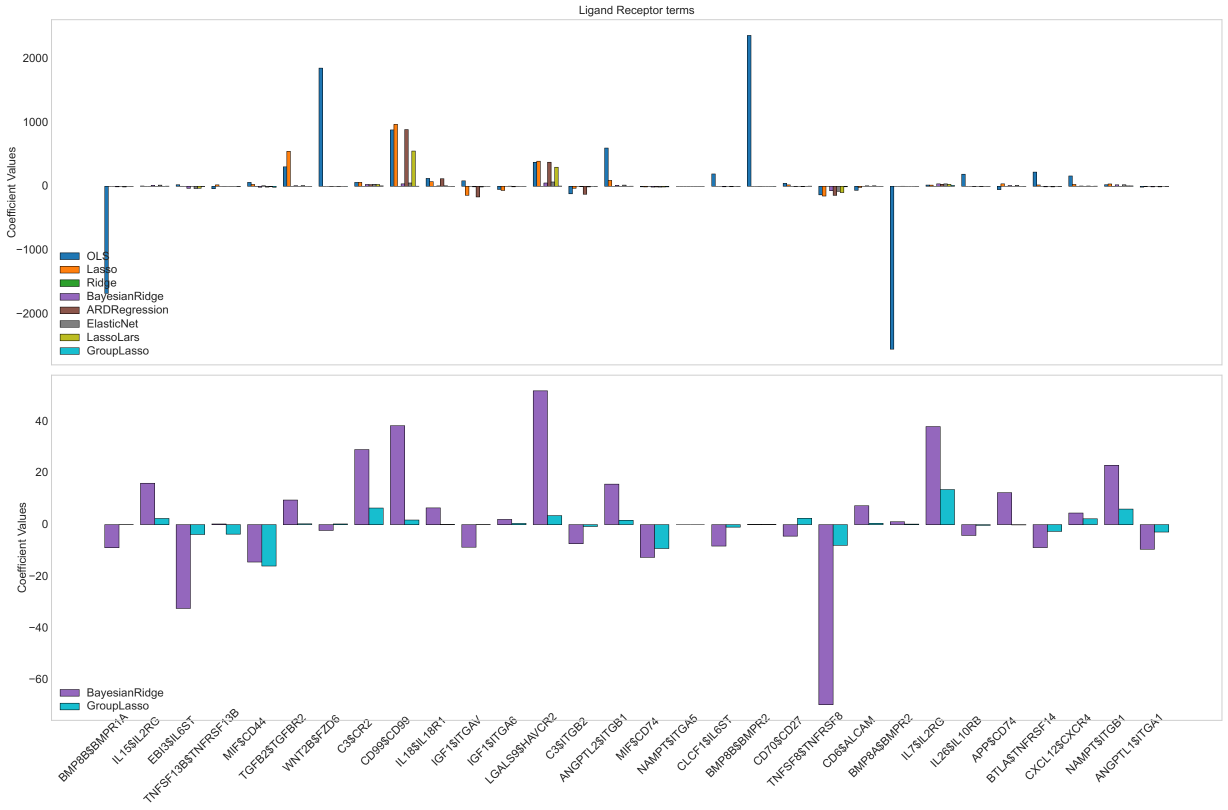
Ligand Receptor terms
Coefficient Values
2000
1000
0
−1000
−2000
OLS
Lasso
Ridge
BayesianRidge
ARDRegression
ElasticNet
LassoLars
GroupLasso
40
20
0
−20
−40
−60
BayesianRidge
GroupLasso
BMP8B$BMPR1A
IL15$IL2RG
EBI3$IL6ST
TNFSF13B$TNFRSF13B
MIF$CD44
TGFB2$TGFBR2
WNT2B$FZD6
C3$CR2
CD99$CD99
IL18$IL18R1
IGF1$ITGAV
IGF1$ITGA6
LGALS9$HAVCR2
C3$ITGB2
ANGPTL2$ITGB1
MIF$CD74
NAMPT$ITGA5
CLCF1$IL6ST
BMP8B$BMPR2
CD70$CD27
TNFSF8$TNFRSF8
CD6$ALCAM
BMP8A$BMPR2
IL7$IL2RG
IL26$IL10RB
APP$CD74
BTLA$TNFRSF14
CXCL12$CXCR4
NAMPT$ITGB1
ANGPTL1$ITGA1

c

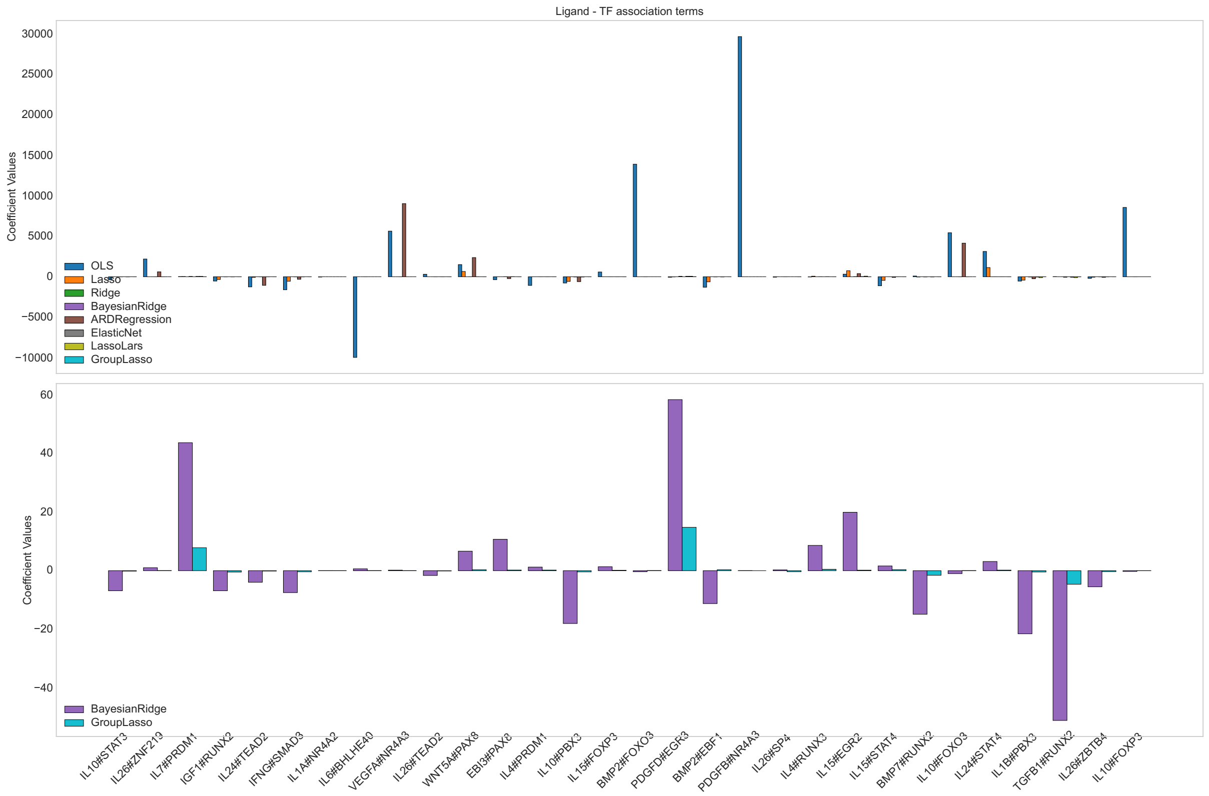
Ligand - TF association terms
Coefficient Values
30000
25000
20000
15000
10000
5000
0
−5000
−10000
OLS
Lasso
Ridge
BayesianRidge
ARDRegression
ElasticNet
LassoLars
GroupLasso
60
40
20
0
−20
−40
BayesianRidge
GroupLasso
IL10#STAT3
IL26#ZNF213
IL7#PRDM1
IGF1#RUNX2
IL24#TEAD2
IFNG#SMAD3
IL1A#NR4A2
IL6#BHLHE40
VEGFA#NR4A3
IL26#TEAD2
WNT5A#PAX8
EBI3#PAX8
IL4#PRDM1
IL10#PBX3
IL15#FOXP3
BMP2#FOXO3
PDGFD#EGR3
BMP2#EBF1
PDGFB#NR4A3
IL26#SP4
IL4#RUNX3
IL15#EGR2
IL15#STAT4
BMP7#RUNX2
IL10#FOXO3
IL24#STAT4
IL1B#PBX3
TGFB1#RUNX2
IL26#ZBTB4
IL10#FOXP3

Comparison of the coefficients estimated by all regression backbones (top) and just Bayesian Ridge and Group Lasso for the three groups of modulators for FOXO1: A. TFs, B. ligand-receptors, C. and ligand-TF terms. TF modulator coefficients remain relatively consistent across models. However, other models, including Bayesian Ridge, estimate extremely high coefficients for interaction terms without group-level penalization, while Group Lasso successfully eliminates uninformative features to reduce false positives.

#### Supplementary Figure 8. Vector plot convergence post FOXO1 KO between Dark Zone and Light Zone GC B cells

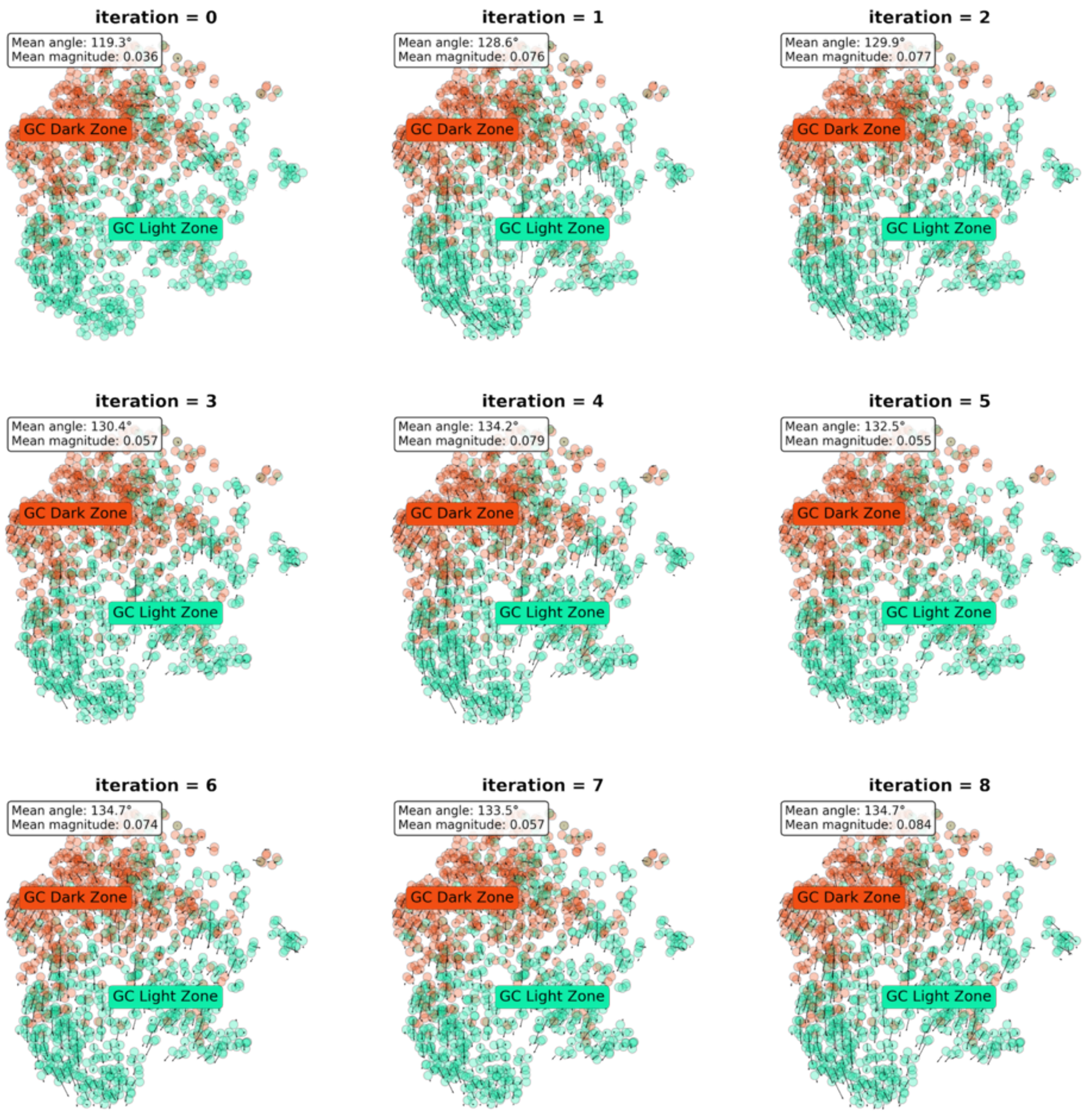


Transition vector plots after 1 through 10 hops for the GC Dark Zone and Light Zone B cells in Slide-Tags Human Tonsil. The transition probabilities converge at approximately 4 hops, which SpaceTravLR uses as its network propagation depth.

Supplementary Figure 9. Vector plot convergence of FOXO1 KO for each cell type

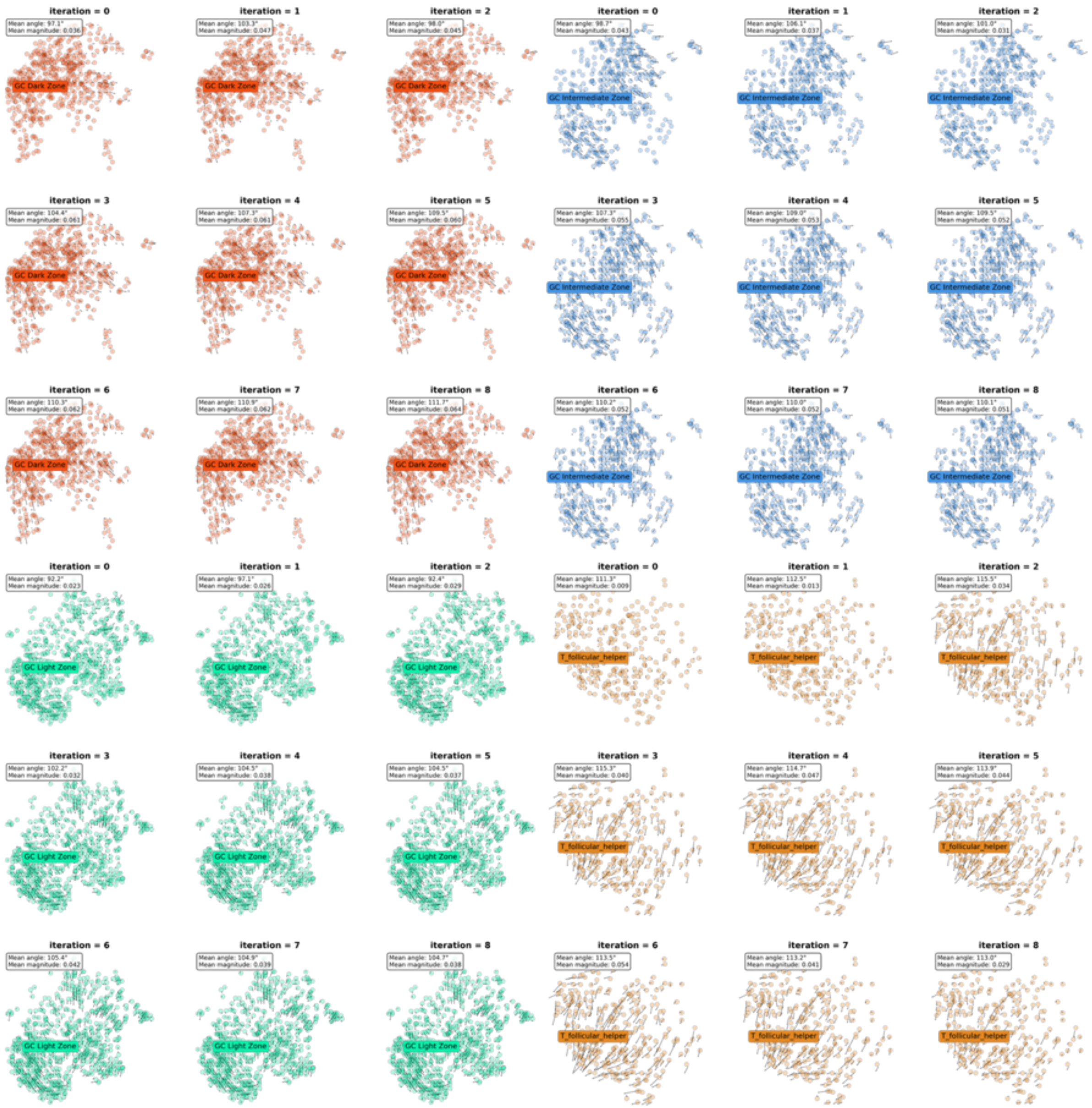

Transition vector plots after 1 through 10 hops for A. GC Dark Zone B cells, B. GC Intermediate/ Marginal Zone GC B cells, C. GC Light Zone B cells, and D. Tfh cells in Slide-Tags Human Tonsil. The transition probabilities converge at approximately 4 hops, which SpaceTravLR uses as its network propagation depth.

#### Supplementary Figure 10: Hypertonic and regulator macrophages in XYZseqV2 Murine Kidney Replicates

a.

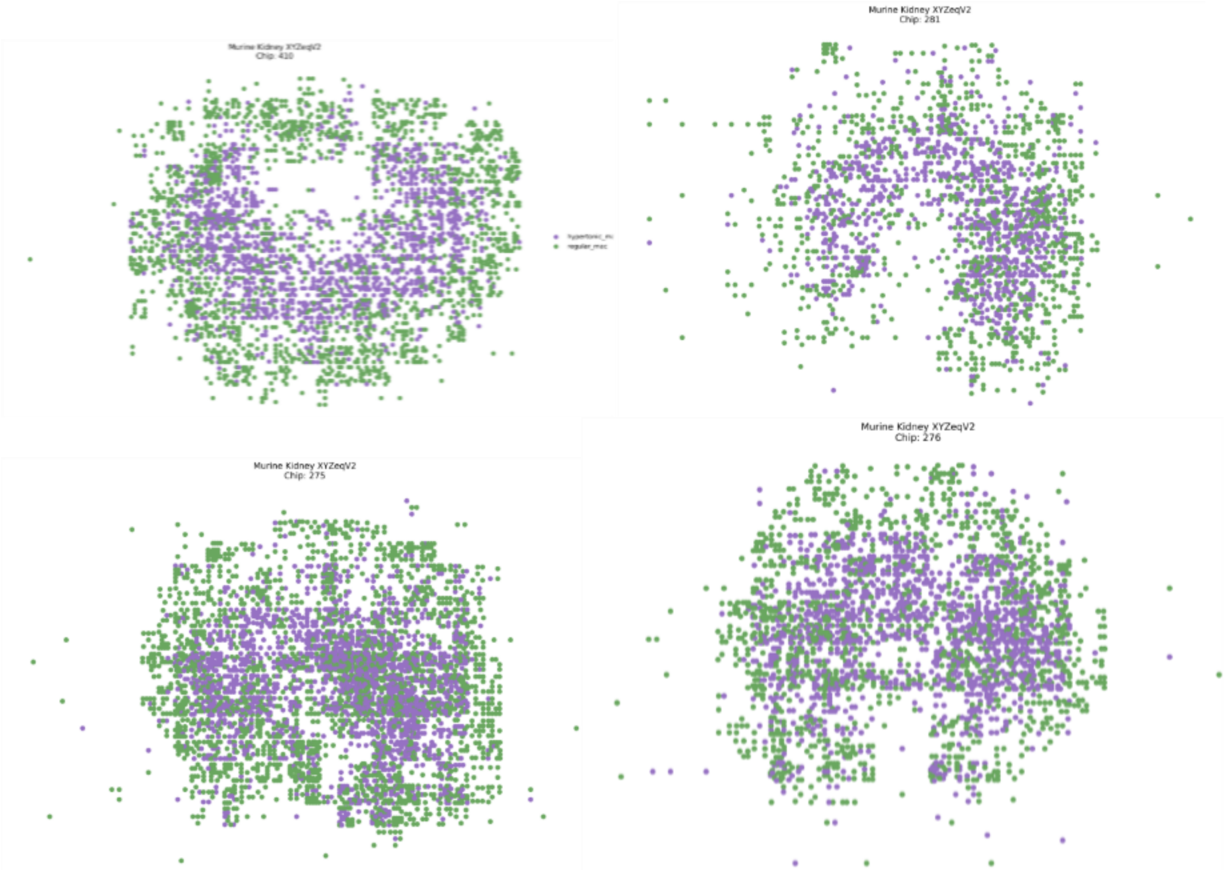


A. 4 separate mouse kidney samples were profiled using the XYZeqv2 pipeline, with clear spatial localization of the two distinct macrophage populations across replicates. Only macrophages are shown here. All samples were processed using the Survey pipeline (https://github.com/survey-genomics/survey).

#### Supplementary Figure 11: Gene-level changes from simulated Tbx6 knockout in SlideSeqV2 mouse embryo

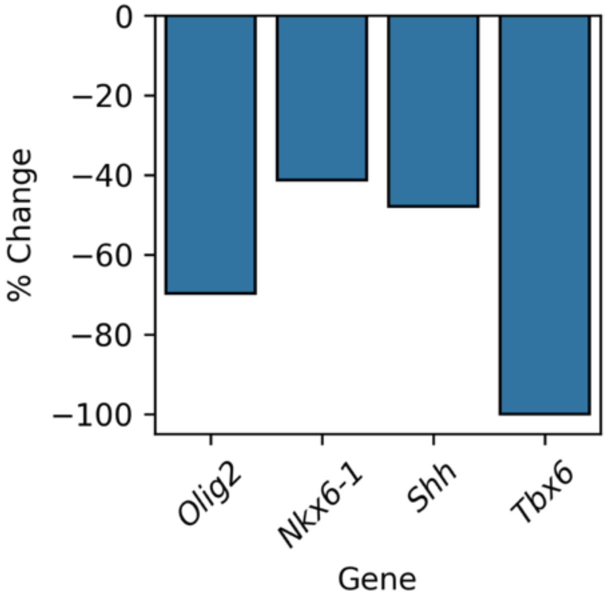


*Tbx6* KO in Slide-seqV2 mouse embryo results in negative percent change of *Olig2*, *Nkx6-1* and *Shh*.

#### Supplementary Figure 12: Additional analyses from gene target screening in Slide-Tags melanoma

a.

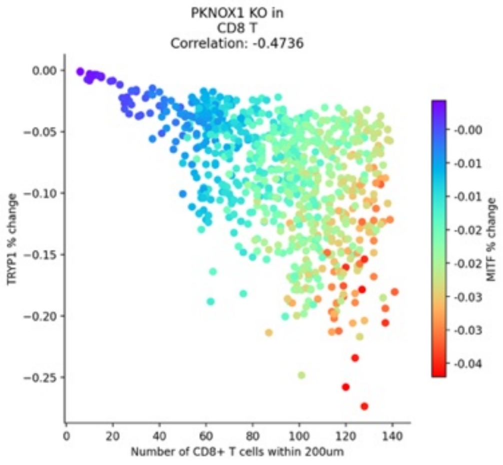


b.

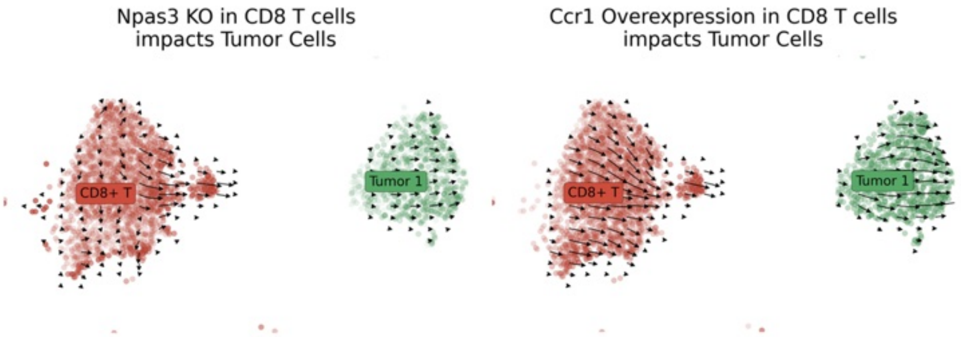


A. In CD8 T cell-specific *Pknox1* overexpression, the percent change of *Typr1* in Slide-tags melanoma tumor cells is negatively correlated with the number of CD8 T cells in their environment. B. Both CD8 T cell specific genetic alterations, *Ccr1* overexpression and *Npas3* knockout, in the Slide-Tags melanoma sample, is predicted to have negative impact on tumorigenicity score.
